# A two-step *in vivo* CRISPR screen unveils pervasive RNA binding protein dependencies for leukemic stem cells and identifies ELAVL1 as a therapeutic target

**DOI:** 10.1101/2022.06.11.494613

**Authors:** Ana Vujovic, Laura P.M.H. de Rooij, Ava Keyvani Chahi, He Tian Chen, Brian Yee, Sampath K Loganathan, Lina Liu, Derek C.H. Chan, Amanda Tajik, Emily Tsao, Steven Moreira, Pratik Joshi, Joshua Xu, Nicholas Wong, Sasan Zandi, Stefan Aigner, John E. Dick, Mark D. Minden, Daniel Schramek, Gene W. Yeo, Kristin J. Hope

**Affiliations:** Department of Biochemistry and Biomedical Sciences, McMaster University, Hamilton, ON, Canada; Department of Medical Biophysics, University of Toronto, Toronto, ON, Canada; Princess Margaret Cancer Centre, University Health Network, Toronto, Ontario, Canada; Department of Cellular and Molecular Medicine, University of California at San Diego, La Jolla, CA, USA; Centre for Molecular and Systems Biology, Lunenfeld-Tanenbaum Research Institute, Mount Sinai Hospital, Toronto, Ontario, Canada; Laboratory Medicine and Pathobiology, University of Toronto, Toronto, ON, Canada; Department of Molecular Genetics, University of Toronto, Toronto, Ontario, Canada; Faculty of Medicine, University of Toronto, Toronto, Ontario, Canada

## Abstract

Acute myeloid leukemia (AML) progression and relapse is fueled by self-renewing leukemic stem cells (LSCs) whose molecular determinants have been difficult to discern from normal hematopoietic stem cells (HSCs) or to uncover in screening approaches focused on general AML cell properties. We have identified a unique set of RNA binding proteins (RBPs) that are enriched in human AML LSCs but repressed in HSCs. Using an *in vivo* two step CRISPR-Cas9-mediated screening approach to specifically score for cancer stem cell functionality, we found 32 RBPs essential for LSC propagation and self-renewal in MLL-AF9 translocated AML. Using knockdown or small molecule approaches we show that targeting key hit RBP ELAVL1 impaired LSC-driven *in vivo* leukemic reconstitution and selectively depleted primitive AML cells vs. normal hematopoietic stem and progenitors. Importantly, knockdown of Elavl1 spared HSCs while significantly reducing LSC numbers across genetically diverse leukemias. Integrative RNA-seq and eCLIP-seq profiling revealed hematopoietic differentiation, RNA splicing and mitochondrial metabolism as key features defining the leukemic ELAVL1-mRNA interactome with the mitochondrial import protein TOMM34 being a direct ELAVL1-stabilized target whose inhibition impairs AML propagation. Altogether, through the use of a stem cell-adapted in vivo CRISPR dropout screening strategy, this work demonstrates that a wide variety of post-transcriptionally acting RBPs are important regulators of LSC-survival and self-renewal and, as exemplified by ELAVL1, highlights their potential as therapeutic targets in AML.

## Introduction

Acute myeloid leukemia is a hematological malignancy characterized by clonal expansion and accumulation of immature myeloid cells. To date, the effective treatment of AML remains a significant unmet clinical need, with standard of care underscored by dismal overall patient survival and high relapse rates^1^. Decades of combined research in the mouse and human contexts have shown that populations of self-renewing leukemic stem cells (LSCs) arising from mutations in hematopoietic stem or progenitor cells are both the seeds of initiation as well as the drivers of progression and relapse in AML^2,3^. Unlike bulk AML, LSCs exhibit unique cellular attributes including quiescence^4,5^, extensive self-renewal, and the capacity to localize to protective microenvironments^6,7^. Because of the critical role LSCs have in propagating AML their effective and specific targeting represents a key therapeutic goal but one that is currently challenged by our poor understanding of the unique molecular drivers of the LSC state. While *in vivo* CRISPR screening is gaining steam as a strategy to identify cancer and leukemia cell dependencies in general^8-10^, to guide the discovery of the highest value therapeutic targets in AML the uniqueness of the cancer stem cell state necessitates novel tailored high throughput screening approaches that can *a priori* identify dependencies not just of progenitors and blasts but of the driving LSCs themselves.

A focus on epigenetic and transcriptional changes that may underlie leukemic behavior has defined a large proportion of investigations into AML-specific targets to date however the extensive post-transcriptional layer as it pertains to LSC function has received comparably little attention. Here, RNA-binding proteins (RBPs) are core effectors, rapidly executing precise control of gene expression by modulating a diversity of RNA properties that include splicing, polyadenylation, localization, stabilization, degradation, and translation. Through association of their RNA-binding domains (RBDs) with consensus sequences in their targets, each RBP can link the fate of many, often functionally related, mRNAs^11^. When dysregulated, RBPs can contribute to disease pathology with a recent genomic study revealing that 50% of RBPs are mutated across a variety of cancer types^12^ and in AML there exist isolated examples of RBPs that have been uncovered as specific pro-LSC factors^13-16^ notwithstanding, in some cases, their necessity for normal HSCs^17-20^. Despite these intriguing cases and the emerging evidence of the importance of RBP-mediated post-transcriptional control in cancer progression^21-25^, this level of regulation has not been systematically explored in AML LSCs. Moreover, with the complement of RBPs on the order of 2,000, these regulators represent a potentially enormous untapped source for therapeutic target discovery.

In addressing these questions, herein we identified a large subset of RBPs selectively enriched in the LSC fraction of AML by interrogating stem and progenitor cell populations in healthy bone marrow (BM) and AML patient samples. We devised a unique two-step serial transplantation *in vivo* CRISPR-Cas9-mediated pooled dropout screen to identify 32 RBPs underlying LSC function and thus of elevated high translational value. These targets, which span a diverse set of RBPs, include the RNA stabilizing factor ELAVL1. We demonstrate the therapeutic potential of targeting LSCs while sparing healthy stem cell counterparts via small molecule inhibition of ELAVL1 in patient-derived AML xenografts and through comprehensive multi-omics profiling of its post-transcriptional regulon reveal mitochondrial metabolism and the mitochondrial protein import regulator, TOMM34, as a key axis through which ELAVL1 sustains LSC function and AML survival.

## Results

### Identification of an RBP subset uniquely enriched in LSCs

To identify a set of RBPs with potential roles in the selective control of human LSCs we performed an expression study of genes encoding proteins encompassed within the human RBP census^26^. Interestingly, as a class RBPs were revealed as significantly enriched in LSCs by Gene Set Enrichment Analysis (GSEA) of a dataset of transcriptionally profiled and functionally validated LSC^+^ and LSC^-^ fractions obtained from 78 AML patients^27^ (Figure 1A, B). We defined the top 500 LSC-enriched RBPs as LSC leading edge (LLE) and evaluated their expression profiles in primitive fractions of human bone marrow (BM) (Supplemental Table 1, Figure 1A, C). Overall, LLE RBPs were expressed at much lower levels in BM HSPCs compared to the entire RBP census (Figure 1C, left panel). Additionally, we identified a subset of RBPs that exhibit uniquely low and/or relatively reduced expression in the long-term (LT)-HSC compartment relative to more committed short-term (ST)-HSCs and multipotent progenitors (MPPs), which we termed LT-HSC low (LHL) RBPs (Figure 1C, right panel). Upon intersecting the LHL and LLE RBPs, we identified a group of 128 RBPs comparatively elevated in LSCs (Figure 1A, D). Because of their low expression in normal LT-HSCs, indicating possibly reduced importance for normal hematopoietic function, we nominated the 128 RBP subset as potentially critical selective LSC determinants with high therapeutic relevance.

**Figure 1.**
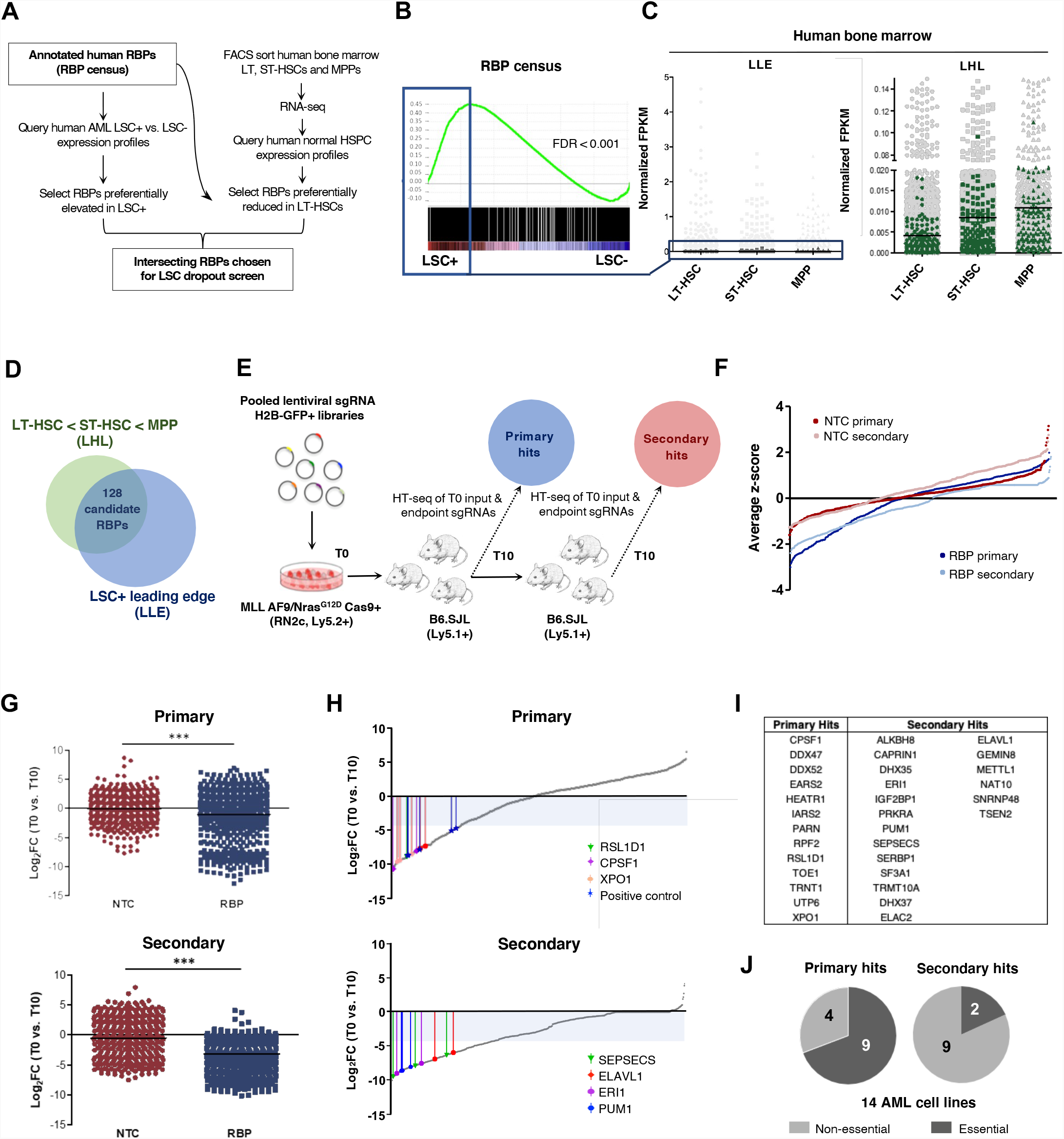
Diverse RBPs are enriched in LSCs and identified as *in vivo* AML LSC essentialities through a two-step pooled *in vivo* CRISPR-Cas9 dropout screen. (A) Overview of *in silico* selection of RBPs preferentially heightened in LSCs and reduced in LT-HSCs. (B) GSEA plot showing LSC-enriched RBPs. The top 500 of the leading edge LSC-enriched RBPs (LLE) are indicated by the blue box. (C) Expression of LSC-enriched RBPs (boxed) in LT-HSC, ST-HSC and MPP populations of human BM (dark grey, left panel) and a subset of RBPs with lowest expression in LT-HSCs of human BM (green, right panel). (D) Selection of the 128 RBPs exhibiting LSC-enriched and HSC-reduced expression. (E) Schematic illustrating the *in vivo* dropout screen. (F) Average ranked dropout z-scores for all sgRNAs in both arms and transplantation rounds. (G) Median log2 fold-change (T10 vs. T0) of unique sgRNAs in the NTC and RBP arms of the screen at the primary (top) and secondary (bottom) endpoints are shown. (H) Median LFC of all sgRNAs within the RBP arm of the screen after the primary (top) and secondary (bottom) rounds. Select top-scoring sgRNAs are indicated with colored bars; shaded area indicates the decreased fold-change of 20 cutoff. (I) RBPs called as hits across primary and secondary screening arms (J) Analysis of general essentiality across a panel of AML cell lines^40^ for the RBPs considered hits at primary endpoints (Primary, left) compared to RBP hits where all targeting sgRNAs dropped out only in secondary recipients (Secondary, right). A gene was considered generally essential if its average log_2_ fold change in abundance was less than -1 in at least 12 of the 14 tested AML lines *in vitro*. n = 2-3 mice per 1° T10 and 2° T0 and T10 replicates. LT-HSC= CD34^+^CD38^-^CD90^+^CD49f^+^; ST-HSC= CD34^+^CD38^-^CD90^+^CD49f^-^; MPP= CD34^+^CD38^-^CD90^-^ CD49f^-^. ***p < 0.001, determined by a two-sided Student’s t test.

### A two-step pooled *in vivo* RBP CRISPR dropout screen in MLL-AF9/Nras^G12D^ leukemia

Primary LSCs are rare^28^, and once isolated, rapidly lost in culture. Consequently, *in vitro* leukemia screens predominately identify genes essential for proliferation (a characteristic property of progenitor cells) but fail to pinpoint genes required for repopulation or self-renewal (hallmark features of *bona fide* stem cells). Therefore, true LSC function can only be assayed *in vivo* by evaluating the capacity of cells to regenerate serially transplantable leukemia^28,29^. To this point we designed a two-step *in vivo* pooled CRISPR-Cas9 screening approach that would identify candidates within the 128 LSC-enriched RBPs capable of regulating the functional property of leukemia reconstitution as measured by primary transplantation. As heightened self-renewal of LSCs contributes to their capacity to serially propagate leukemia, we reasoned that a secondary transplantation step would identify RBPs that uniquely control LSC self-renewal (Figure 1E). We selected a Cas9-expressing MLL-AF9/NRas^G12D^ mouse leukemia (RN2c) as our *in vivo* system^30^, on the basis of its immunophenotypically well-defined and relatively abundant LSC fraction, validated surrogacy of human MLL-AF9 counterpart disease, including its therapeutic targets and its heightened expression of the candidate RBPs to be screened as compared to healthy mouse hematopoietic stem and progenitor cells (HSPCs)^29,31-34^ (Supplemental Figure 1A). We designed lentiviral sgRNA constructs ^30,35^ targeting all 128 LSC-enriched RBPs (4-5 sgRNAs per RBP) wherein annotated early exon RNA binding motifs were targeted where possible to encourage maximum negative selection. Positive control sgRNAs that impair *in vivo* RN2c cell fitness^30^, were combined with >400 non-targeting control (NTC) sgRNAs in a separate arm of the screen (Supplementary Table 2). RN2c cells were infected with either the NTC or RBP targeting library pools and serially transplanted into recipient mice, with sgRNA representation captured post-infection and at each transplant endpoint (Figure 1E).

Over the course of transplantation, transduced H2B-GFP^+^ fractions in the RBP-targeting arm gradually decreased, whereas in the NTC arm the H2B-GFP levels remained stable (Supplemental Figure 1B), findings paralleled at the level of sgRNA representation (Figure 1F, Supplemental Figure 1C,D. When quantitatively assessed using the MAGeCK algorithm^36^, the median log2 fold-change (LFC) of sgRNA abundance as compared to that on the day of transplant was significantly lower at the end of the primary transplant compared to that on the day of transplant in the RBP arm, but not in the NTC arm, and this further increased in magnitude following secondary screening, indicating selective and progressive loss of RBP-targeting sgRNAs (Figure 1G, Supplemental Figure 1E).

### Classification of primary and secondary depleting sgRNAs and target RBPs

To identify RBPs important for leukemic repopulation and LSC-driven propagation, we set a stringent threshold where at least 2 of its targeting sgRNAs must dropout with a fold change greater than 20, a threshold reached for all positive controls tested (Supplemental Figure 1F). Using this selection criterion, we identified 32 hit RBPs, with 13 RBPs reaching our threshold in primary recipients (“primary hits”) and 19 achieving the 2 sgRNA depletion threshold only upon passage through secondary recipients (“secondary hits”) (Figure 1H, I). GO annotations indicate that the screen hits are involved in diverse RNA metabolic processes/interactions and molecular pathways including nitrogen compound metabolism, tRNA processing, ribosome biogenesis and mRNA processing, indicating dependence on a broad range of post-transcriptional regulation (Supplemental Figure 1G, H). The most significant hit RBPs for which the greatest number of sgRNAs dropped out included RSL1D1, CPSF1 (a mRNA cleavage and polyadenylation factor) and XPO1 (mediator of RNA nuclear export) in primary transplants, and ELAVL1, SEPSECS1, ERI1 and PUM1 as significant dropouts only in the secondary round, each distinguished by diverse roles in RNA control with functions ranging from mRNA stabilization, tRNA:selenocysteine synthesis, histone mRNA degradation and translational repression respectively (Figure 1H). Of note, XPO1 and PUM1 have been identified as drivers in human AML LSCs and mouse leukemia respectively^37-39^, supporting our screen’s capacity to identify *bona fide* LSC regulators.

We next analyzed all screen hit RBPs against a previously reported Gene Essentiality analysis of genome-scale *in vitro* CRISPR-Cas9 screening of a panel of 14 AML cell lines. Notably, 69.2% (9/13) of primary hits were found to be generally critical for leukemia cell line propagation while only 42.0% (8/19) secondary hit RBPs were found to be critical for growth across the majority of lines^40^. Importantly when considering those secondary hits in which all sgRNA dropouts called occurred only within secondary recipients, essentiality *in vitro* was found for only 18.2% (2/11) (Figure 1J, Supplemental Figure 1I). Altogether our results suggest that serial *in vivo* screening can effectively identify LSC-drivers including genes regulating LSC repopulation and self-renewal that appear underrepresented or dispensable in *in vitro* dependency screens. Moreover, our results showcase the novel insights that can be gained through the use of the two-step screening approach where the unique secondary screening arm we have employed served to uncover a host of effects masked in the primary transplant arm. Given the critical contribution of self-renewal in LSC-serial reconstitution, this secondary screening arm thus has a high likelihood of having captured *bona fide* LSC-specific events.

### Validation of individual sgRNAs identifies ELAVL1 as a top-scoring RBP

To validate the outcome of our two-step *in vivo* drop out screen we selected top-scoring primary and secondary hit RBPs for individual knockout in RN2c and following their independent repression we assessed the effects in vitro, compared the *in vivo* growth dynamics to that observed in the screen, and quantified the LSC compartment within the leukemic grafts. For each sgRNA used, we verified highly efficient CRISPR-induced indel formation at the targeted locus^41^ (Supplemental Figure S2A). Next, we evaluated *in vivo* propagation of RN2c transduced with these individual sgRNAs in comparison to a non-essential control sgRNA targeting Ano9^30^ (Figure 2A). In line with the screen results, we observed that sgRNAs targeting the primary hits (CPSF1 and RSL1D1) strongly depleted in the first round of transplantation, while sgRNAs targeting secondary hits (ELAVL1, SEPSECS and ERI1) showed more moderate depletion of H2B-GFP^+^ cells in the primary transplantation followed by a strong depletion upon secondary transplantation (Figure 2B, C). These results confirmed that individual knockout events replicated their dynamics as tracked in the pooled screen setting. We next assessed the effects of these sgRNAs in cultured RN2c cells and found that in all cases the knockout of our hit RBPs exhibited either negligible or less detrimental effects on *in vitro* growth in comparison to those observed in the long-term *in vivo* context (Supplemental Figure S2B). Lastly, we quantified the effect of individual gene depletions on the LSC-enriched cKit^+^ fraction^31,34^ within detectable H2B-GFP^+^ grafts. We observed a relative decrease of this fraction for all sgRNAs tested supporting an LSC-specific impairment of RN2c cells *in vivo* upon genetic ablation of hit RBPs (Figure 2D). The approach of considering both canonical and non-canonical RBPs, selecting candidates based on selectively elevated expression patterns in LSC and pairing this with screening in an *in vivo* serial transplantation setting thus allowed us to uncover unique regulators of LSCs highlighting the potential functional and clinical relevance of RBPs in AML.

**Figure 2.**
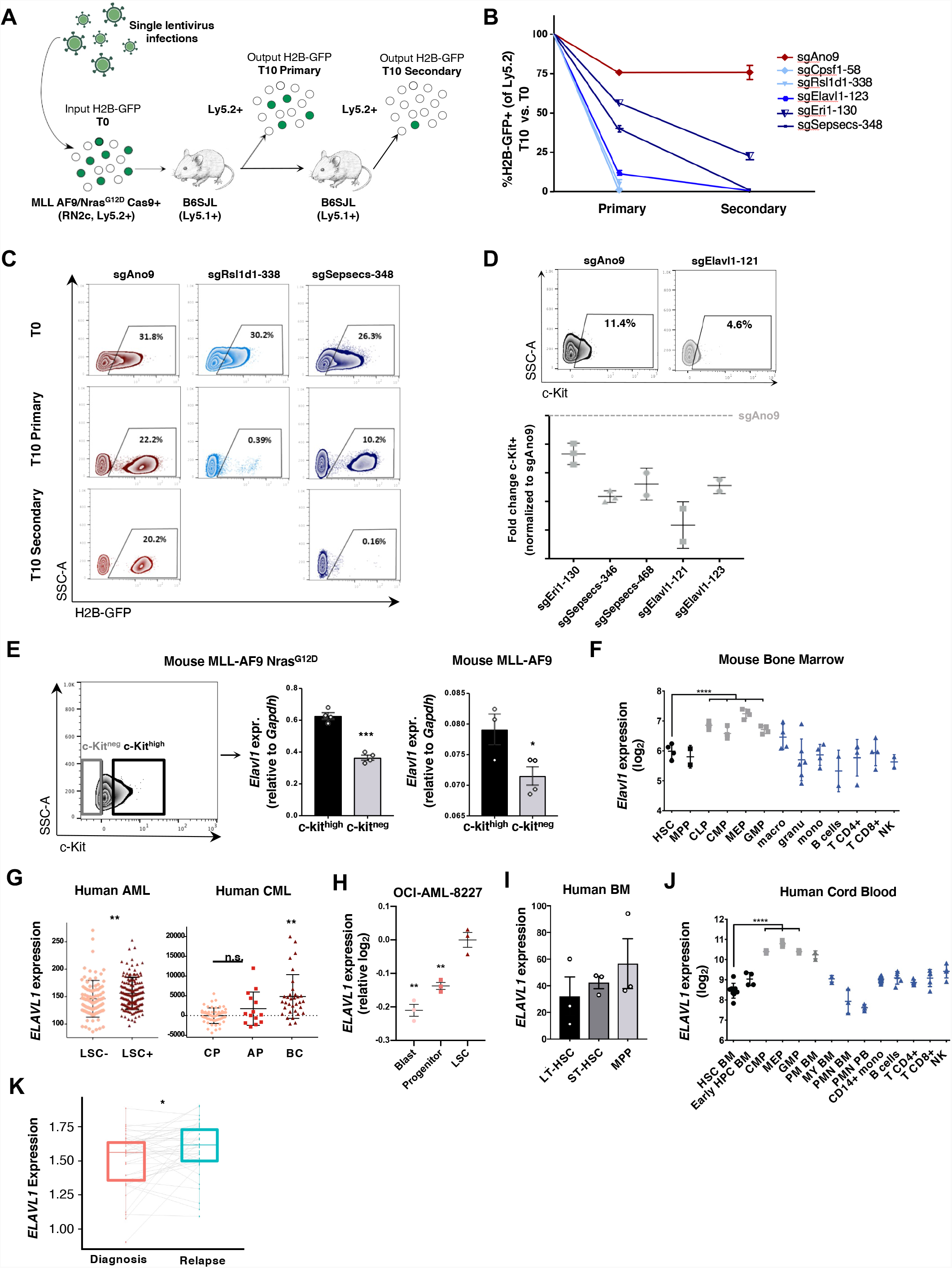
Independent hit knockout validation replicates pooled CRISPR-Cas9 screen dropout dynamics and identifies ELAVL1 as a top LSC dependency. (A) Schematic illustrating the in vivo screen validation strategy. (B) Percentage of H2B-GFP+ cells in the output graft as compared to the T0 input. sgAno9 is the negative control and darkening shades of blue correlate with increasing time in vivo before sgRNA dropout. (C) Representative flow plots of RN2c cells sampled at each time point (T0, T10 primary, T10 secondary) are shown. (D) Fold change (FC) of cKit+ fractions within H2B-GFP+ populations of BM samples with >5% H2BGFP+ of Ly5.2+. (E) Transcript expression of Elavl1 in LSC+ (cKithigh, top 25%) and LSC-(cKitneg) MLL-AF9 NrasG12D (RN2c) cells (left panel; flow sorting gates are shown) and MLLAF9 cells (right panel). (F) Levels of Elavl1 mRNA across various subpopulations of the mouse hematopoietic hierarchy, adapted from BloodSpot. Mature hematopoietic lineages, progenitor populations and primitive populations are shown in blue, grey and black, respectively. (G) Human ELAVL1 expression in LSC+ and LSC-AML subfractions27 and throughout CML disease stages (adapted from Radich et al. 2006 and www.oncomine.org) are shown in left and right panels, respectively. (H) Expression of ELAVL1 in subpopulations of OCI-AML-8227 cell line. (I) ELAVL1 expression levels in highly purified long term (LT)-HSC (CD34+CD38-CD90+CD49f+), short-term (ST)-HSC (CD34+CD38-CD90+CD49f-) and multipotent progenitors (MPP; CD34+CD38-CD90-CD49f-) from human bone marrow. (J) Transcript levels of ELAVL1 across sorted subfractions of the human cord blood hematopoietic hierarchy, adapted from BloodSpot43. Mature hematopoietic lineages, progenitor populations and primitive populations are shown in blue, grey and black, respectively. Normalized fragments per kilobase million (FPKM) are shown. (K) ELAVL1 expression levels across paired diagnosis-relapse primary human AML samples. *p < 0.05, **p < 0.01, ***p < 0.001, ****p < 0.0001, as determined by a two-sided Student’s t test.

From all of the top-scoring secondary hit RBPs independently validated, the sgRNA with the strongest depletion of cKit^+^ cells over the course of *in vivo* leukemic propagation was one that targets Elav-like protein 1 (Elavl1), an RBP primarily characterized for its role in regulating gene expression via stabilization of its RNA targets (Figure 2D). Additionally, while deletion of Elavl1 resulted in pronounced *in vivo* depletion of RN2c, a more moderate growth inhibition was observed *in vitro* (Supplemental Figure S2B). Likewise, in the human AML cell line THP-1, which also possesses MLL-AF9/NRas^G12D^ mutations, we found that CRISPR-mediated knockout of ELAVL1 did not significantly increase apoptosis and yielded only a modest ∼20% reduction in cell growth and progenitor CFU output (Supplemental Figure 2C-G), together implicating Elavl1 as an LSC regulator that may be underprioritized by an *in vitro* screening approach. Furthermore, we found that the LSC-enriched cKit^high^ expressing fraction of RN2c cells showed significantly elevated transcript levels of *Elavl1* compared to cKit^neg^ cells (Figure 2E, left and middle panel). This was also observed in the cKit^high^ and cKit^low^ fractions of a separately derived MLL-AF9 mouse model devoid of Ras mutations (Figure 2E, right panel, Supplemental Figure 2H-J), indicating an LSC-specific increased expression of this RBP in MLL-AF9-driven leukemias. We next examined the expression of Elavl1 in publicly available datasets of mouse HSPCs^42,43^. Compared to MPPs, Elavl1 is decreased at the protein level in the HSC fraction of normal mouse BM (Supplemental Figure 2K). Moreover, *Elavl1* shows a progressive elevation in expression within mouse HSPCs peaking in downstream megakaryocyte-erythroid progenitors (MEP), indeed suggesting that its heightened expression in the stem cell context could be unique to AML (Figure 2F).

Human and mouse ELAVL1 show >90% protein sequence conservation^44^, and consistent with having passed the expression filters necessitated by the screen’s candidate selection strategy, *ELAVL1* transcripts are significantly enriched in human AML LSC^+^ fractions^27^ (Figure 2G, left panel). When analyzed in a dataset of 91 BCR-ABL-driven chronic myeloid leukemia (CML) patient samples, a gradual increasing pattern is observed for *ELAVL1* transcript levels across the progressively more aggressive chronic to blast crisis (BC) phases with the highest levels observed in BC (Figure 2G, right panel), the most LSC-enriched phase^45^. Additionally, the expression of ELAVL1 in a primary AML model system propagated in vitro but possessing a defined functional hierarchy (OCI-AML-8227) demonstrates a significant enrichment in the most primitive population of LSCs compared to downstream clonogenic progenitors and terminal blasts^46^ (Figure 2H). Importantly, consistent with the mouse system, *ELAVL1* levels are lower in human HSCs and show a step-wise increase throughout the human HSPC hierarchy peaking in MEPs (Figure 2I,J). Moreover, when *ELAVL1* is evaluated in the bulk cells across a cohort of primary AML samples representing 27 distinct categories of cytogenetic abnormalities, it is in almost every case heightened in expression in leukemic blasts relative to normal HSC counterparts^47-52^ (Supplemental Figure 3L), indicating that its elevation in leukemia is common and largely agnostic to underlying genetic abnormalities. Above median levels of *ELAVL1* expression did not show any significant prognostic trend in two independent cohorts of AML patients^52,53^, however in a pooled set of expression profiles from 39 paired AML diagnosis-relapse samples^3,54,55^ *ELAVL1* expression is significantly increased upon relapse (Figure 2K). Our *in vivo* screening results considered together with these robust expression profiles in normal and leukemic samples are strongly predictive that ELAVL1 could be an important cross cutting driver of LSC function across leukemic subtypes with potentially reduced dependence in healthy HSCs.

### Depletion of Elavl1 expression selectively impairs the murine LSC compartment

Given the potentially selective importance of Elavl1 in LSCs we next took an RNAi-based approach to pursue loss-of-function studies in multiple genetically diverse LSCs and normal HSPCs. Using our lentiviral systems, we knocked down Elavl1 in both non-Ras mutated MLL-AF9 and BC-CML mouse leukemias (Figure 3A, Supplemental Table 2). We observed significantly decreased colony-forming ability in shElavl1-transduced MLL-AF9 cells relative to shLuciferase controls (Figure 3B, Supplemental Figure 3A). Upon transplantation of infected MLL-AF9 BM cells we observed a significant loss of transduced Ametrine^+^ populations within shElavl1-transduced secondary grafts suggesting that leukemic cells, and LSCs in particular, are sensitive to reduced Elavl1 (Figure 3C). Furthermore, cell surface marker analysis revealed that, in contrast to controls, the LSC-enriched cKit^high^ fraction^32,33^ of shElavl1-infected populations specifically decreased over the course of *in vivo* propagation (Figure 3D). When repeated in the distinct LSC-driven BC-CML mouse model, that has been used to dissect clinically relevant insights into the corresponding human disease (Supplemental Figure 3B-E)^56-58^, we again observed impaired serial *in vivo* leukemic reconstitution upon knockdown of Elavl1 (Supplemental Figure 3F). Moreover, shLuciferase and shElavl1-transduced BC-CML LSCs revealed increased levels of apoptosis upon Elavl1 knockdown (Supplemental Figure 3G, H). These findings demonstrate that in genetically distinct types of mouse myeloid leukemia, Elavl1 repression has significant inhibitory effects on LSC-mediated *in vivo* leukemic growth.

**Figure 3.**
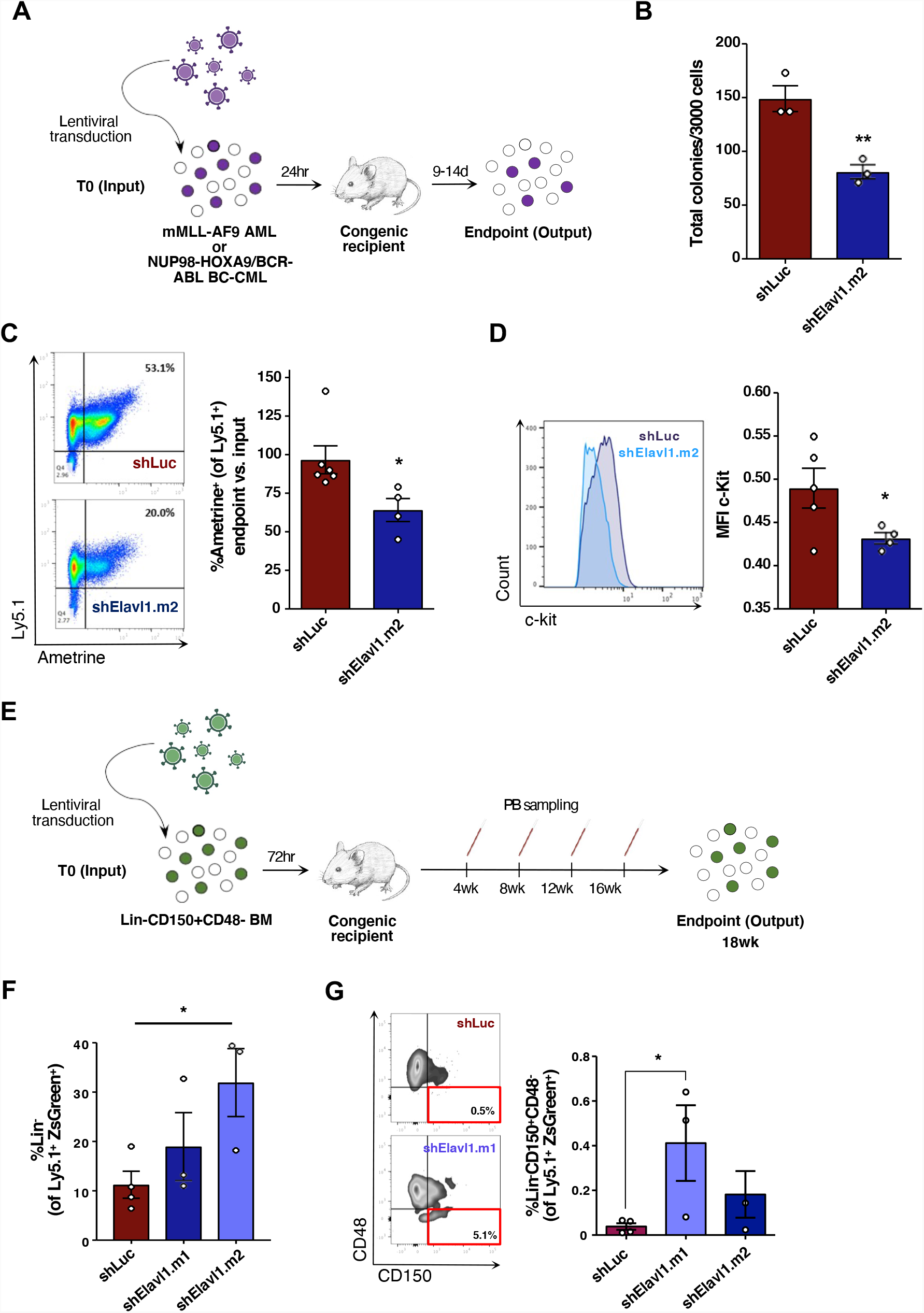
Elavl1 knockdown impairs in vivo leukemic propagation and spares healthy LTHSCs. (A) Schematic illustrating Elavl1 loss-of-function in vivo transplantation assays in mMLL-AF9 and BC-CML mouse models. (B) CFU output from shLuciferase-and shElavl1-infected MLL-AF9 mouse leukemic BM, 10 days post-plating (n= 3). (C, D) Flow cytometric analysis of the normalized output vs. input Ametrine+ fractions (C) and c-Kit mean fluorescence intensity (D) of shLuciferase-and shElavl1-infected MLL-AF9 BM at the secondary transplant endpoint. Representative flow plots and histograms are shown at left in each panel. (E) Schematic of in vivo evaluation of Elavl1 knockdown in normal mouse bone marrow stem and progenitor cells. (F, G) At 18 weeks post-transplant, Ly5.1+ BM from all recipient mice was harvested and the Lin-(F) and Lin-CD150+CD48-(G) fractions quantified by flow cytometry. *p < 0.05, determined by a two-sided Student’s t test.

We next wanted to assess the effect of a similar level of Elavl1 repression on normal stem cell function, as ideal anti-leukemic therapeutics should spare healthy HSCs. Previous reports present contrasting conclusions on the effects of Elavl1 loss in primitive murine hematopoietic cells. In a conditional knockout model, inducible deletion of Elavl1 in native mice did not alter bone marrow LSK percentages on short term follow up^59^, whereas a separate study suggested LSK Flt3^+^CD34^-^ cells are reduced in the 12 week grafts derived from primitive bone marrow cells transduced with a single hairpin directed against Elavl1^60^. Methodological differences and the possibility for off-target effects in the shRNA study complicate the interpretation of these disparate findings. In addition, Yilmaz *et al* have reported that inclusion of SLAM family markers, which were not explored in the latter study, markedly increase the purity of HSCs from reconstituted mice^61^. To address this, we lentivirally delivered shRNAs (ZsGreen^+^) into Lin^-^CD150^+^CD48^-^ mouse HSCs (Ly5.1^+^) and assayed their function over long-term hematopoiesis by competitive transplantation (Figure 3E). At 4 weeks post-transplant, during which early multipotent progenitors are the dominant contributors to reconstitution levels, the shElavl1-infected grafts in the peripheral blood showed a significant decrease of 2.3-2.7-fold compared to control. After this point however, changes to ZsGreen^+^ proportions in shElavl1 grafts normalized and tracked closely to control grafts (Supplemental Figure 3I). Most importantly, at the 18-week endpoint, the mean percentage of Lin^-^ and the highly HSC-enriched Lin^-^CD150^+^CD48^-^ fraction within the shElavl1 BM grafts were either not reduced or elevated relative to shLuciferase (Figure 3F, G). In addition, we observed no lineage skewing within shElavl1 grafts (Supplemental Figure 3J). Together our findings suggest that while constitutive repression of ELAVL1 impairs an early population of progenitors, HSCs that are essential for long-term reconstitution are relatively maintained.

### ELAVL1 knockdown promotes myeloid differentiation and inhibits *in vivo* leukemic propagation in human AML

Given the profound effects of Elavl1 loss on LSCs from genetically diverse murine leukemia models, we next evaluated the functional role of ELAVL1 in human AML by introducing shScramble-(control) and shELAVL1-expressing lentiviruses into primary AML cells (Supplemental Figure 4A, Supplemental Table 2, 3). Immunophenotyping of infected cells by flow cytometric analysis revealed increased myeloid populations, as measured by CD14^+^ and CD11b^+^ expression (Figure 4A), as well as an overall enrichment of these antigens as demonstrated by their increased median fluorescence intensities (MFI) compared to control (Supplemental Figure 4B). We also observed increased cell death with ELAVL1 reduction as measured by 7AAD^+^ populations (Figure 4B). Colony forming unit assays established post-infection revealed decreased total AML-CFU outputs in ELAVL1 knockdown conditions compared to the control (Supplemental Figure 4C). Together, these results demonstrate that ELAVL1 plays an important role in regulating leukemic proliferation and differentiation.

**Figure 4.**
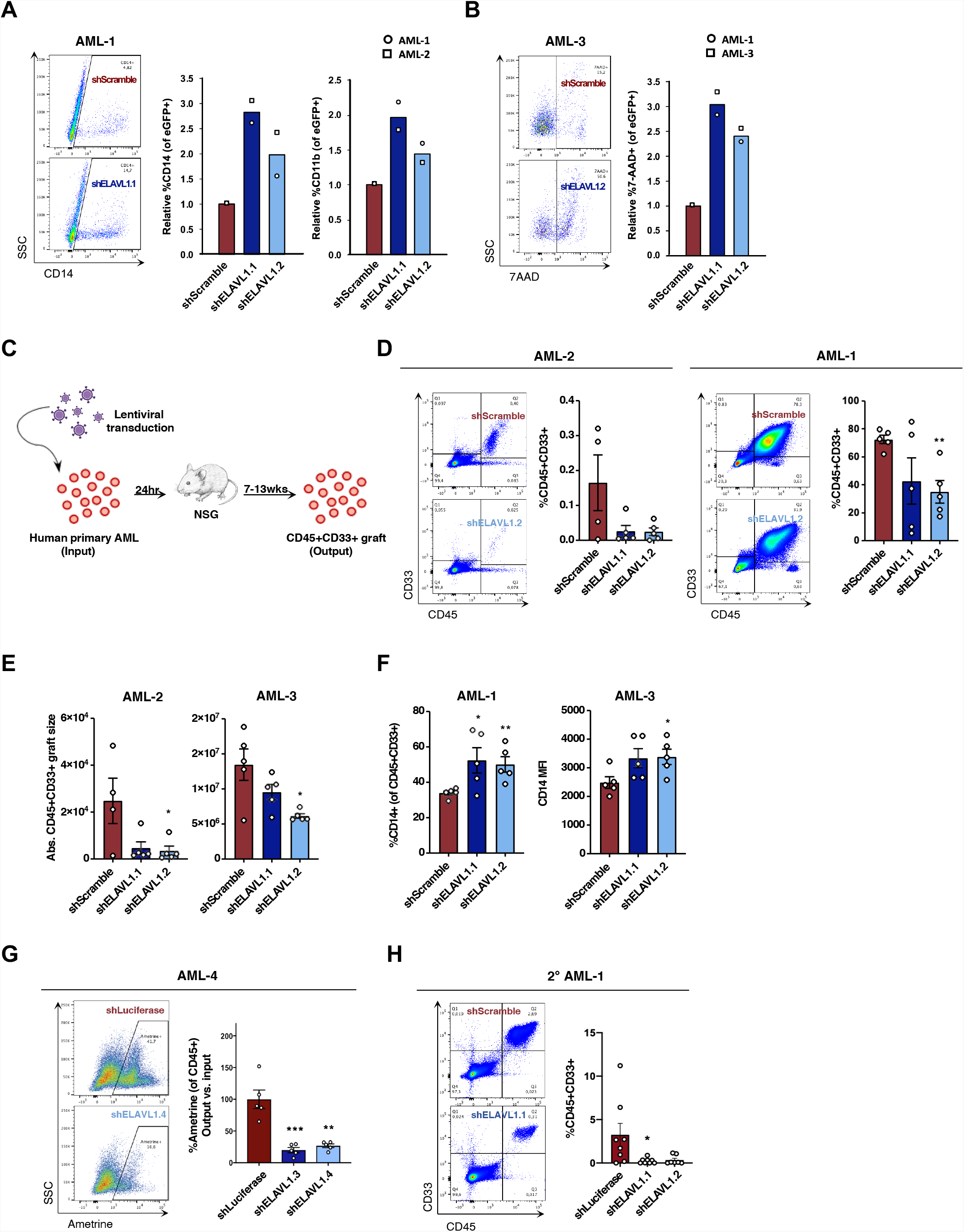
ELAVL1 knockdown impairs in vivo leukemic engraftment. (A, B) Flow cytometric evaluation of (A) CD14+ and CD11b+ and (B) 7AAD+ fractions of shScramble-and shELAVL1-infected primary AML cultures 10 and 2 days post-infection, respectively. (C) Schematic illustrating in vivo ELAVL1 loss-of-function leukemic repopulation assays. (D) Quantitative analysis of shELAVL1-infected primary AML cells at endpoint showing %CD45+CD33+ grafts in BM and (E) absolute graft size based on total cell counts in femurs and tibiae of recipient mice. (F) Flow cytometric analysis of CD14+ populations within CD45+CD33+ grafts in right femur and bone marrow at endpoint. (G) Analysis of Ametrine+ fractions of shLuciferase-or shELAVL1-infected) fractions within the injected right femur CD45+ grafts at endpoint. (H) Flow cytometric analysis of leukemic grafts in the bone marrow of secondarily transplanted recipient mice. Representative flow plots are shown. *p < 0.05, **p < 0.01, ***p < (?), determined by a two-sided Student’s t-test.

To directly assess the role of ELAVL1 in LSC-driven malignant propagation, we performed xenotransplantation assays using shELAVL1-infected unsorted primary AML specimens and captured the effects of ELAVL1 knockdown by measuring leukemic engraftment (CD45^+^CD33^+^) in recipient mice at endpoint (Figure 4C). We tested three separate patient samples and in all cases knockdown of ELAVL1 significantly impaired leukemic growth *in vivo* as demonstrated by an ∼2-fold reduction in the percentage and total number of human AML cells (Figure 4D, E). Immunophenotyping of leukemic grafts depleted of ELAVL1 also uncovered an elevation of CD14^+^ (both in percentage and MFI) and CD11b^+^ populations (Figure 4F; Supplemental Figure 4D). In a fourth sample where intermediate-level infection was achieved post-transduction, the infected (Ametrine^+^) populations were tracked from the day of transplant (input) to endpoint (output) (Figure 4C, Supplemental Figure 4E). At 12 weeks post-transplant we observed a significant ∼4-fold decrease in leukemic engraftment in ELAVL1 depleted AML recipient mice relative to control (Figure 4G). To test the effects of ELAVL1 loss on LSC function, we performed the gold standard assay of serially transplanting the bone marrow from primary xenografts into secondary recipients. At the 6-week endpoint, we indeed observed a further reduction of leukemic burden in the bone marrow of recipients of shELAVL1-relative to shScramble-transduced primary graft cells (Figure 4H). Together, this data indicates that, in contrast to the more modest growth defects of ELAVL1 loss on cultured cells from the human AML THP-1 cell line described above (Supplemental Figure 2C-G), its repression in primary AML not only substantially compromises leukemic growth *in vivo* but directly impairs LSC activity and thus long-term leukemic reconstitution.

### Inhibition of ELAVL1-RNA interactions selectively impairs AML at the LSC level

To investigate the effects of small molecule inhibition of ELAVL1 in AML cells, we used the compound Dihydrotanshinone-I (DHTS), reported to inhibit the interaction of ELAVL1 with its mRNA targets^62^. In an initial set of experiments, THP-1 cells treated with DHTS showed trends toward increased apoptosis and myeloid maturation as indicated by elevated CD14^+^ populations as well as decreased cell division as determined by PKH26 labelling (Supplemental Figure 5A-D). We next tested the impact of DHTS on primary AML cells *in vitro*. Immunophenotyping analysis of DHTS-treated cultures demonstrated significantly increased MFIs of both mature myeloid antigens, CD14 and CD11b, as well as CD33 48 hours post-DHTS treatment compared to control conditions (Figure 5A, B; Supplemental Figure 5E). At the same timepoint, we observed a significant increase in cell death as measured by AnnexinV^+^7AAD^+^ staining (Figure 5C). CFU assays carried out in the presence of DHTS yielded significantly lower malignant myeloid progenitor colonies ranging from 2 to 3-fold compared to vehicle control, indicating leukemic progenitors were effectively compromised (Figure 5D, Supplemental Figure 5F). To confirm the cell-selective context of this drug, we performed CFU assays on normal HSPC populations from lineage depleted human umbilical cord blood (CB) cells. Here, the total progenitor activity remained unaltered with decreases observed only in BFU-E colonies, indicating a general insensitivity of normal committed progenitors to DHTS (Figure 5E). Finally, to assess the functional consequences of ELAVL1 inhibition by DHTS, we treated two primary AML samples and assessed their xenotransplantation potential. Evaluating AML engraftment at 9 weeks post-transplant we observed an impairment in leukemic growth in recipient mouse peripheral blood and bone marrow and a trend towards elevated CD14^+^ populations in the residual leukemic bone marrow graft (Figure 5F, Supplemental Figure 5G).

**Figure 5.**
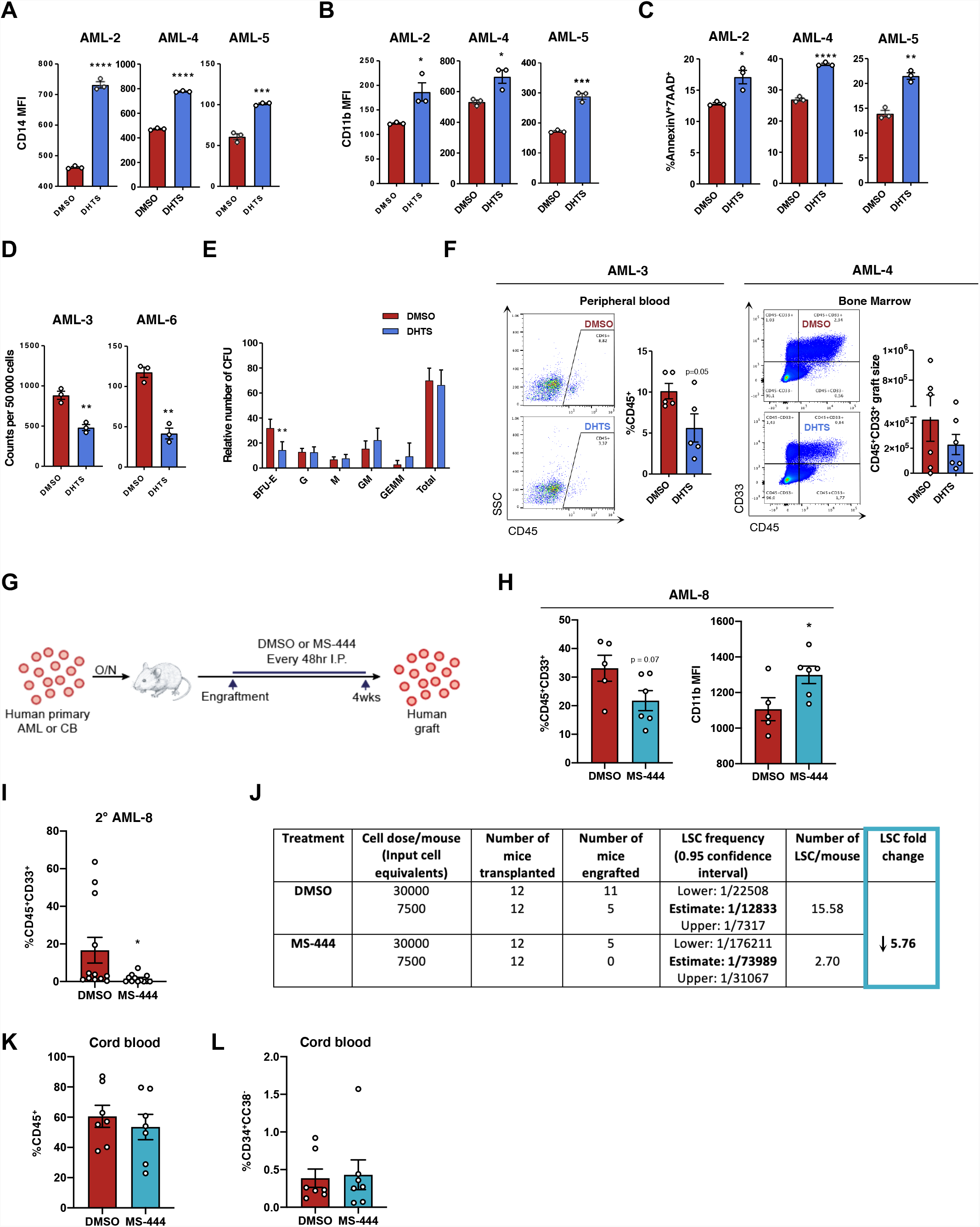
Small molecule-inhibition of ELAVL1 differentially targets leukemia-propagating vs healthy hematopoietic cells. (A-C) Flow cytometric evaluation of the median fluorescence intensities of CD14 (A) and CD11b (B) and percentage of dead cells (AnnexinV^+^7AAD^+^) (C) in human primary AML treated for 48hr with DMSO or DHTS (5.4uM). n=3 replicate experiments. (D, E) CFU output from primary AML (D) or lineage depleted CB cells (E) treated with DMSO or DHTS (1.1 μM). n = 2 independent CB units assessed over two independent experiments, n=4 replicates for each condition. (F) Flow cytometric analysis of leukemic grafts in peripheral blood at 9wks post-transplant and the CD45+CD33+ graft size based on total cell counts in bone marrow at 8wks post-transplant. Representative flow plots are shown. (G) Schematic illustrating *in vivo* administration of DMSO or MS444 in human primary AML engrafted mice. (H) Quantitative analysis of engraftment levels (left) and CD11b expression (in the CD11b+CD45+CD33+ fraction, right) in the bone marrow of human primary AML engrafted mice treated with DMSO or MS-444. (I, J) Leukemic engraftment levels (I) and limiting dilution analysis (J) of secondary recipients transplanted with bone marrow from primary mice treated with DMSO-or MS-444. (K, L) Flow cytometric analysis of hematopoietic engraftment (K) and the primitive (CD34+CD38) HSC population (of the CD45+ graft) (L) in CB-transplanted mice treated with DMSO or MS-444. n.s. = not significant, *p < 0.05, **p < 0.01, ***p < 0.001, ****p < 0.0001 determined by a two-sided Student’s t test.

To test the therapeutic potential of targeting ELAVL1 we used a bioavailable and more potent inhibitor of ELAVL1-mRNA target binding, MS-444, for in vivo administration over the course of primary AML leukemic reconstitution assays. Upon engraftment, mice were treated with 20mg/kg of MS-444 or vehicle control (I.P.) every 48hr for 4 weeks (Figure 5G). At endpoint, the bone marrow from the MS-444-treated recipients showed decreased leukemic burden and significantly increased myeloid differentiation as measured by CD14 expression (Figure 5H). To assess the effects of MS-444-driven inhibition of ELAVL1 on LSCs, we serially transplanted the bone marrow from primary mice into secondary recipients in two doses. At endpoint, the mice that received the MS-444-treated bone marrow demonstrated a dramatic impairment of leukemic reconstitution in which half of the mice did not have a leukemic graft at all (Figure 5I). In addition, using limiting dilution analysis to evaluate LSC frequency differences by virtue of binary engraftment calls in low vs high cell dose-transplanted recipients, we determined that LSCs decreased in MS-444 treated mice by 5.8-fold in comparison to vehicle treated controls (Figure 5J). Together, this demonstrates that ELAVL1 inhibition by MS-444 significantly impairs the ability of LSCs to drive long-term leukemic propagation.

Finally, to evaluate the effects of MS-444 on normal tissues and primitive normal hematopoietic cells in particular, we treated human umbilical cord blood (CB)-engrafted mice with the same in vivo regimen (Figure 5G). Here, we observed no evidence of toxicity with all animals being healthy and showing no evidence of adverse reactions throughout the entirety of the treatment regimen, as was observed in our MS-444-treated leukemic xenografts. Importantly, the CB grafts of MS-444-treated recipients showed no change in their levels compared to vehicle-treated mice and the HSPC (CD34+) and most primitive HSC-enriched populations (CD34+CD38-) were also preserved at control levels in the grafts of MS-444-treated recipients (Figure 5K, L, Supplemental Figure 5H). Moreover, MS-444-treated grafts were normal in all respects and exhibited no evidence of lineage skewing (Supplemental Figure 5I). Together, this data demonstrates not only the potential of small molecule inhibition of ELAVL1 to treat AML but the selectivity of this strategy in targeting the LSC population known to drive long-term propagation of the leukemia.

### Elavl1 enacts LSC-supportive post-transcriptional circuitry

Given our findings that ELAVL1 is essential for LSC maintenance and its known role as a stabilizer of its RNA targets, we sought to comprehensively examine ELAVL1’s underlying mechanism using global transcriptomic profiling upon its knockout in the LSC-rich RN2c cells. Elavl1 knockout resulted in 243 upregulated and 47 downregulated transcripts (Figure 6A; Supplemental Table 4). To capture the full spectrum of coordinated changes in functionally related processes influenced by ELAVL1 loss, we performed GSEA. Consistent with the functional impairment of LSCs in our ELAVL1-depleted primary AML xenografts, we uncovered 252 enriched gene sets and identified that in particular, signatures of LSC maintenance are diminished while those of myeloid differentiation are activated with Elavl1 loss (Supplemental Figure 6A). Enrichment mapping of Elavl1-dependent transcriptional changes furthermore revealed the most significantly negatively regulated clusters relate to metabolism, cytoplasmic translation, and mitochondrial respiration (Figure 6B; Supplemental Table 5). The outcome of our transcriptome profiling thus highlights the possibility that the role of ELAVL1 in LSC maintenance involves reprogramming of certain cellular metabolic processes.

**Figure 6.**
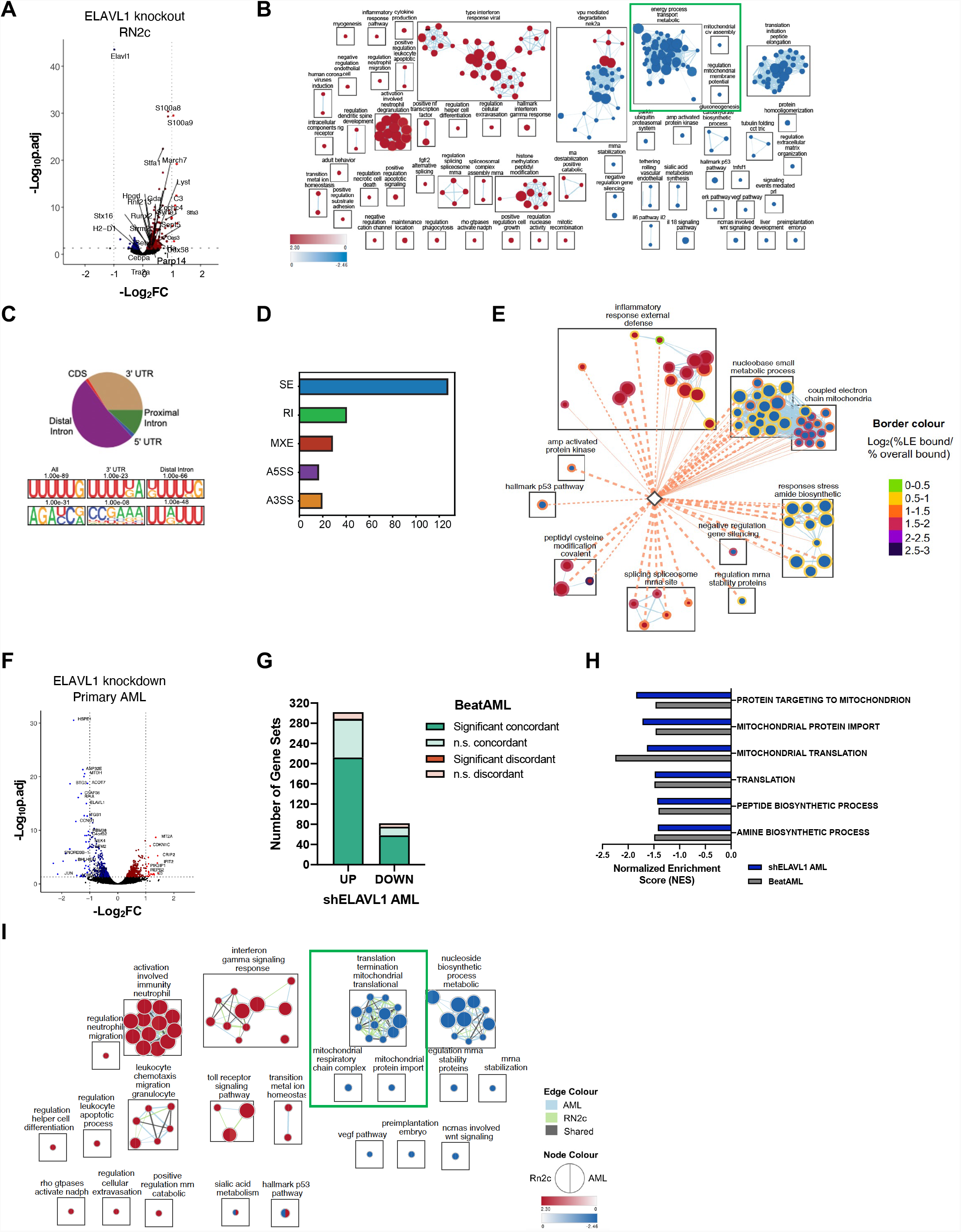
Characterization of the ELAVL1-dependent circuitry in primitive leukemic cells. (A) Volcano plot of differential gene expression in Elavl1-knockout RN2c RNA-seq. Genes with significant differences in expression are highlighted. Blue and red dots represent genes significantly down or up respectively using a padj<0.05 (RNA-seq) cutoff. (B) Enrichment map of genesets significantly enriched (FDR<0.1) in the transcriptome of Elavl1-knockout RN2c cells. (C) Distribution of Elavl1 eCLIP peaks in different genic regions (top) and most common ELAVL1-binding motif sequences (bottom). (D) Distribution of splicing events in Elavl1-knockout RN2c cells. (E) Enrichment map of pathways enriched (FDR<0.1) in the Elavl1-knockout RN2c transcriptome and containing >5% of leading-edge transcripts bound by Elavl1. Colour of borders is based on enrichment of transcript binding to leading edge relative to geneset background. (F) Volcano plot of differential gene expression in ELAVL1-KD human primary AML. Blue and red dots represent genes significantly down or up respectively using a padj<0.05 cutoff. (G) Number of pathways in the human ELAVL1-knockdown AML transcriptome that are significantly or non-significantly concordant and discordant in the BeatAML RNA-seq data set. (I)Normalized enrichment scores (NES) of downregulated mitochondrial gene sets in the human ELAVL1-knockdown RNA-seq data set (highlighted by the green box in Supplemental Figure 6I).Enrichment map of gene sets significantly (FDR<0.25) altered in both ELAVL1-knockdown human AML and Elavl1-knockout RN2c transcriptomes.

We next sought to identify direct LSC-specific binding targets of ELAVL1 by performing enhanced cross-linking immunoprecipitation followed by deep-sequencing (eCLIP-seq)^63^. To enable efficient eCLIP pulldown, we used mouse BC-CML cells, which similarly to RN2c are enriched in Elavl1-dependent LSCs (Supplemental Figure 3B-F), while importantly exhibiting the requisite high expression of ELAVL1 protein (Supplemental Figure 6B). We discovered 4345 significant ELAVL1-binding peaks across 1548 genes with a preference for intronic regions, suggesting that ELAVL1 can act in the nucleus on pre-mRNA species, as well as 3’UTRs, consistent with typical binding profiles underlying its role in mRNA stabilization (Figure 6C, Supplemental Table 6), In agreement with this, we identified nuclear and cytoplasmic localization of ELAVL1 in both mouse leukemic BM and human primary AML cells (Supplemental Figure 6C). In line with previous findings^64,65^, an enrichment for U-rich binding motifs was identified within regions bound by ELAVL1 (Figure 6C). Lastly, biological processes overrepresented among Elavl1-bound transcripts included hematopoietic differentiation, transcriptional control, mRNA processing and mRNA splice regulation (Supplemental Table 6).

To identify transcripts directly regulated by ELAVL1, we interrogated the expression outcomes of ELAVL1-bound transcripts from the eCLIP-seq in the ELAVL1-depleted transcriptome. From this, we identified 156 genes whose significant differential transcript levels appears directly linked to physical association with ELAVL1 (Supplemental Figure 6D, Supplemental Table 6). These genes were predominantly upregulated in response to Elavl1 loss (Supplemental Figure 6D), a significant trend found amongst all bound transcripts (Supplemental Figure 6E). Within the bound and downregulated transcripts upon Elavl1 loss were several LSC enforcers and oncogenes including *Gpr56, Dazap1* and the previously appreciated direct target of ELAVL1, *Myc*^66-71^ (Supplemental Figure 6D). In contrast, the direct ELAVL1 targets upregulated upon its loss include *Neat1*, a differentiation promoter and a known ELAVL1 target^72,73^, as well as a significant overrepresentation of mRNA splicing regulators (Supplemental Figure 6D, F, Supplemental Table 6). Noting this latter class of targets as well as the pre-mRNA binding action of ELAVL1 (Figure 6C, Supplemental Figure 6F), we explored mRNA splicing changes upon Elavl1 depletion. This analysis revealed 230 diverse changes with exon skipping being the most common event (Figure 6D, Supplemental Table 7). Interestingly, among these exon skipping events, we again identified dysregulated mitochondrial genes. More specifically, we noted the presence of several altered exon splicing events (delta percent spliced in (PSI) > 0.25) in genes with characterized roles in mitochondrial function and integrity including *Ewsr1, Samd8, Nsun3, Pam16* and *Mrtfl1* (Supplemental Figure 6G).

Given that our multi-omics analyses implicate mitochondrial regulation as a probable hub downstream of ELAVL1, we examined whether ELAVL1 binding contributes to direct regulation of mitochondrial genes. Overall, we found that ELAV1 is bound to leading edge transcripts in 85% of the significantly enriched gene sets observed in the ELAVL1 knockout transcriptome, including mRNA splicing, activated immune functions and metabolic processes (Figure 6E). We noted that an average of 9.3% (range: 5.7-15%) of the leading-edge transcripts in 14 gene sets related to the electron transport chain are also bound by ELAVL1, a significant 1.6-fold enrichment over the gene set background (t-test p < 0.05) (Figure 6E).

Altogether, profiling of the ELAVL1-directed regulon via integrated analyses of the RNA-interactome and transcriptome strongly implicates mitochondrial activity as a critical axis through which ELAVL1 supports LSCs.

### ELAVL1 repression impairs leukemic mitochondrial function

We next aimed to profile ELAVL1 dependencies integral to the maintenance of human AML. To this end, we performed RNA sequencing in primary patient AML cells upon ELAVL1 knockdown and identified 333 upregulated and 376 downregulated genes (Figure 6F, Supplemental Table 8). To assess the clinical relevance of these ELAVL1-associated programs, we compared the shELAVL1 transcriptome to RNA-seq data of 494 bulk AML samples from the BeatAML clinical dataset^74^. We observed a positive correlation in overall gene expression profiles of below-median ELAVL1 expressers and ELAVL1-knocked down AML, particularly among differentially expressed genes (ρ = 0.32, p<10e-10). Assessing pathway-level changes in the shELAVL1 transcriptome, we again observed dysregulation of LSC maintenance and myeloid differentiation gene signatures (Supplemental Figure 6H, Supplemental Table 8), demonstrating ELAVL1’s role in maintaining stemness features in human AML. Overall, we discovered 410 aberrantly regulated pathways of which 70% are significantly concordant with the BeatAML dataset with no discordant events (Figure 6G, Supplemental Figure 6I). Most importantly, echoing our findings in the LSC-enriched RN2c setting, even when profiled at the bulk level, we observed a negative enrichment of mitochondrial import and translation in AML samples with experimentally or disease-specific reduced levels of ELAVL1 (Figure 6H, Supplemental Figure 6I). The strong mirroring of these and other expression signatures suggests that ELAVL1-dependent programming actively shapes the transcriptomic landscape of clinical AML.

Finally, in an effort to uncover core ELAVL1-mediated biological processes underlying its support of human LSC, we compared pathway-level changes in ELAVL1-depleted bulk human AML and murine LSC-rich RN2c cells. Within commonly altered gene sets we again confirmed the recurring theme of activation of myeloid differentiation programs, and repression of constituents of mitochondrial function and integrity (Figure 6I). Given that dependence on mitochondrial function has recently emerged as a selective regulatory mechanism of the LSC compartment^75^, we hypothesized that ELAVL1’s essential role in LSC may be mediated by its control over mitochondrial processes. Indeed, flow cytometric measurements of mitochondrial activity via MitoTracker dye staining showed significant decreases in both murine RN2c and human primary AML cells upon ELAVL1 depletion (Figure 7A,B, Supplemental Figure 7A,B). These results, in combination with our comprehensive molecular profiling support that maintenance of mitochondrial activity is an important mechanism through which ELAVL1 achieves its critical and selective role in supporting AML LSCs.

**Figure 7.**
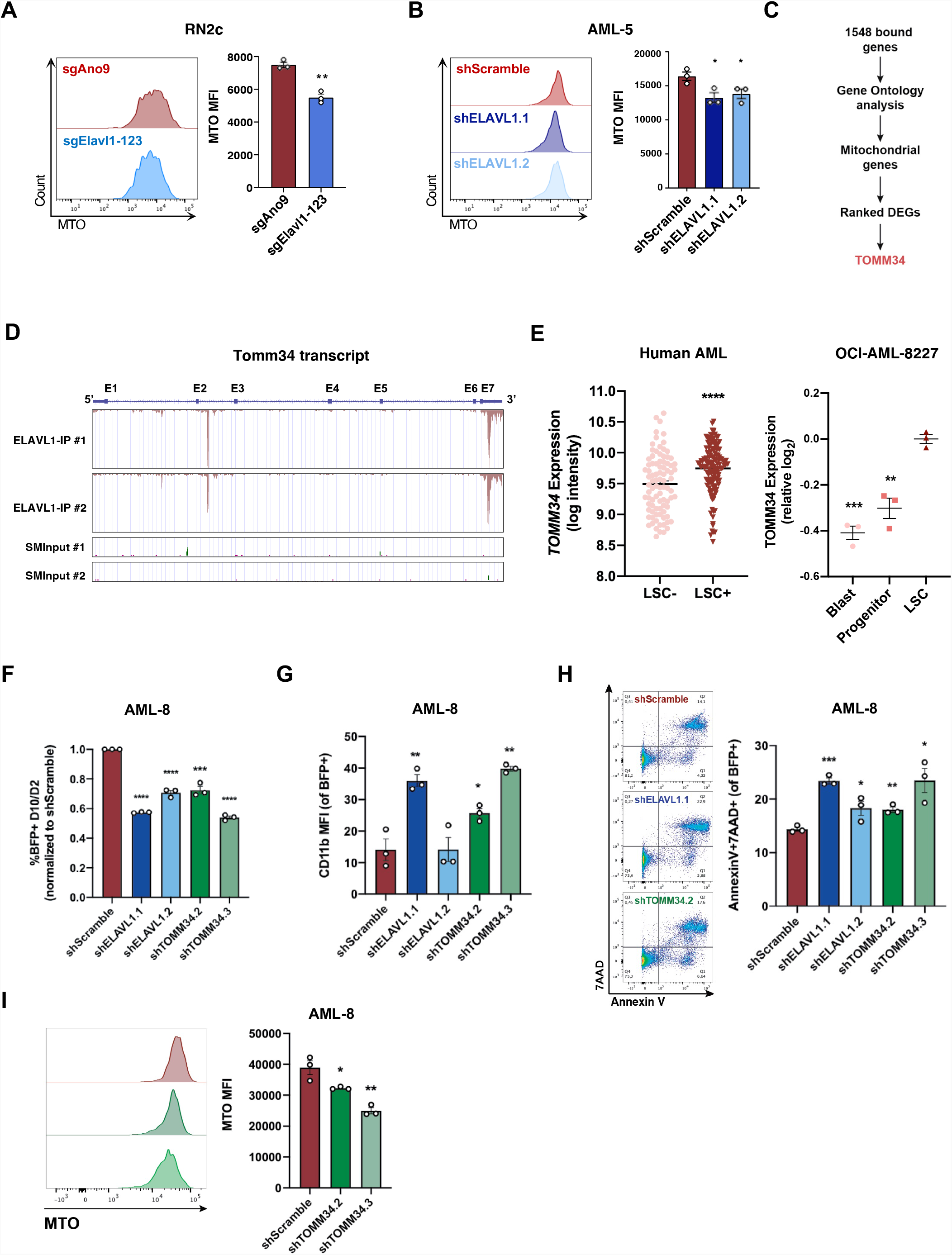
TOMM34 is a direct effector of ELAVL1, essential for mitochondrial metabolism and maintenance of primitive AML cells. (A, B) Quantification of MitoTracker Orange (MTO) median fluorescence intensities in RN2c (A) and human AML cells (B) following ELAVL1 depletion. (C) Flow chart illustrating the steps in identifying a top downregulated mitochondrial gene directly bound and regulated by ELAVL1. (D) UCSC Genome Browser tracks showing ELAVL1-binding peaks along the Tomm34 transcript in reference to size-matched small input (SMInput) controls. (E) Expression of TOMM34 in human AML LSC+/LSC-fractions (left)^27^ and the OCI-AML-8227 cell line (right). (F-H) Flow cytometric analysis of proliferation (BFP+) (F), myeloid differentiation (G) and cell death (H) in ELAVL1 and TOMM34 depleted human primary AML compared to controls. (I) MTO analysis of TOMM34 depleted human primary AML.

### TOMM34 is a direct effector of ELAVL1 in AML cells

Given that our integrative bioinformatic analyses and functional experiments demonstrate that mitochondrial control is a key axis through which ELAVL1 drives its phenotype in AML, we sought to identify an ELAVL1-direct effector that might underlie this control. To do this, we returned to the list of 1548 transcripts bound by ELAVL1 and using Gene Ontology analysis isolated mitochondrial genes and ranked them based on their significance of differential expression in the human AML RNA-seq data set (Figure 7C). This analysis identified translocase of outer mitochondrial membrane 34 (TOMM34) as the ELAVL1-directly bound mitochondrial gene with the topmost downregulation upon ELAVL1 loss (Figure 7C,D). TOMM34 is a co-chaperone that facilitates the heat shock protein HSP70/HSP90-mediated import of mitochondrial preproteins^76-78^ and while it has been associated with poor survival in solid tumors^79,80^, its physiological role in mitochondrial metabolism and involvement in leukemia pathogenesis is unknown. Interestingly however, as for ELAVL1, we find *TOMM34* transcripts are significantly enriched in the LSC^+^ fraction of human AML (Ng et al.) with preferential expression at the protein level in the LSC compartment of the patient-derived 8227 AML model (Figure 7E). Moreover, *TOMM34* transcript levels also correlates with *ELAVL1* expression in the diverse BeatAML data set of primary AML samples, consistent with the predicted role of ELAVL1 in stabilizing TOMM34 by virtue of ELAVL1’s association with its 3’UTR (Supplemental Figure 7C).

To determine the functional similarity of TOMM34 to ELAVL1, we performed immunophenotyping and apoptosis assays upon shRNA-mediated TOMM34 depletion in human primary AML in parallel to ELAVL1 knockdown. Here, we observed that repression of TOMM34 significantly impairs proliferation and promotes myeloid maturation and cell death, mirroring closely the phenotype observed by ELAVL1 loss (Figure 7F-H, Supplemental Figure 7D). Finally, to assess the effects of TOMM34 loss on mitochondrial function, we used MitoTracker Orange in TOMM34-depleted human primary AML cells and show that, as we observed upon ELAVL1 loss, mitochondrial activity is indeed significantly compromised compared to controls (Figure 7I). Altogether, this data indicates that TOMM34 is a positively regulated direct downstream target of ELAVL1 through which it enacts its role in maintaining mitochondrial function for AML survival.

## Discussion

Elucidation of LSC-targeted therapies remains a profound unmet need for AML, a cancer that under standard of care therapy is characterized by high rates of relapse and poor long-term survival. Current efforts to uncover key molecular LSC dependencies have largely overlooked the potentially target-rich class of post-transcriptional regulators. *In vivo* functional genetic screens have been important in showcasing the insights that come from interrogating cancer cell dependencies in the setting of a complex, in situ niche however have yet to be applied in such a way as to uncover cancer stem cell-specific regulators. We describe the first LSC-focused pooled CRISPR dropout screen in AML performed uniquely in the *in vivo* serial transplantation setting. This approach has uncovered regulators of clear importance to LSCs and thus may serve as a novel strategy the field can capitalize on to prioritize candidates of maximum clinical interest more systematically. Moreover, with a focus on post-transcriptional regulators as an understudied class of candidate LSC determinants, this unique approach identified RBPs essential to the repopulating and/or self-renewal function of LSCs. Of the 128 RBPs preferentially expressed in LSCs that we systematically screened, 32 were required for *in vivo* leukemic propagation. This identification of a large number of RBPs underlying LSC function combined with the enrichment in expression of the entire class of RBPs in LSCs speaks to an intriguing dependence of LSCs on RBP-driven post-transcriptional control mechanisms that warrant future in-depth investigations into their therapeutic potential. This list of RBPs is diverse and encompasses regulators known to influence virtually all aspects of RNA metabolism. In addition, our screen has highlighted specific cellular pathways known to be under the control of certain hit RBPs, which have been previously implicated in LSCs (i.e. rRNA metabolism^81^), and others including tRNA modification and ribosome biogenesis, not before appreciated for their specific contribution to AML LSC function. Given the parallels between LSCs and other tissue-specific cancer stem cells^2^ our findings portend value in exploring the extent to which RBP-mediated regulation contributes to the functionality of cells driving diverse cancer types.

Of the identified RBP hits in our screen, ELAVL1 imparted amongst the most significant inhibitory effects to LSCs and leukemic cell growth over the course of the screen and when independently targeted was essential for mouse and human leukemic reconstitution. Overexpression of ELAVL1 has been reported in various solid tumor types^82-84^, as well as myeloid and lymphoid leukemias^45,85,86^. However, its role in the stem cell compartment of leukemia has thus far not been addressed. Our functional evaluation in combination with transcriptional profiling following ELAVL1 knockout in LSC-enriched cells indicates that ELAVL1 enforces a larger molecular profile that maintains LSC stemness, restricts differentiation and preserves survival. Importantly, we find that ELAVL1 disruption via shRNA or small molecule interventions significantly impair LSC self-renewal but allow for a relative sparing of the compartment of normal mouse and human HSPCs, raising the significance of this RBP as a candidate therapeutic target in AML. Our results with DHTS and MS-444 further suggest that inhibition of RBP-mRNA interactions might also be as effective as depletion of the RBP itself, a concept also hinted at by the Pfam targeting strategy of our screen. As many non-canonical RBPs have functions beyond regulation of RNA^87,88^, explicit reliance on their RNA-binding features for LSC function may also provide novel opportunities for anti-leukemic intervention. Indeed, this concept is exemplified by our results with MS-444 administered *in vivo*, which demonstrate the clear therapeutic potential and relevance of targeting ELAVL1-mRNA interactions to directly impair LSC function and reduce leukemic burden.

ELAVL1 has well-described roles in stabilizing pro-cell growth mRNAs through association with AU-rich regions within 3’UTRs^65,89-92^. While ELAVL1 has also been described to influence splicing in non-hematopoietic tissues^93-95^, as RBPs generally confer distinct cell context specific effects on the global splicing status, the nature of any splicing regulation by ELAVL1 in AML LSCs has not been known^96^. In LSCs we find that the majority of ELAVL1’s binding events are intronic, in line with the findings of Mukherjee *et al*., who documented a previously unrecognized importance for intronic ELAVL1 binding in pre-mRNA stabilization^65^. In addition, we reveal that 15 ELAVL1-bound transcripts appear dependent on ELAVL1 for stabilization as compared to 141 whose expression is elevated upon ELAVL1 deletion. Among the latter ELAVL1-bound and negatively regulated transcripts we also observed an enrichment in regulators of splicing. Together these findings not only indicate that a much larger pool of total bound transcripts exist for which ELAVL1 interaction may be repressive in these cells, but that this non-canonical role of ELAVL1 is selective to splicing regulators. Consistent with this, we found that global splicing was indeed significantly altered upon ELAVL1 knockout, which supports a critical role for ELAVL1 in propagating a specific LSC alternative splicing (AS) program both directly and indirectly. These findings are intriguing given the burgeoning appreciation for AS dysregulation as a result of splicing regulator mutations that promote leukemic transformation, propagation and relapse^97,98-100^ and the potential for altered expression of splicing regulators to promote pathogenic splicing in other cancers^101-103^.

Our integrative omics analysis uncovered an ELAVL1-nucleated post-transcriptional circuitry in LSCs that in large part coalesces on a signature of oxidative phosphorylation conservation. In addition to being a previously uncharacterized target of ELAVL1, this finding is particularly interesting in light of studies that situate mitochondrial metabolism as a critical axis that LSCs are selectively dependent on relative to normal HSCs. More specifically, AML and CML LSCs maintain a decreased spare reserve glycolytic capacity in comparison to HSCs rendering them especially vulnerable to inhibitory strategies that target mitochondrial metabolism. Such strategies including targeting of mitochondrial protein synthesis, mitochondrial DNA replication, amino acid metabolism or mitochondrial protein degradation, selectively kill LSCs while sparing HSCs^104-106^. Intriguingly, even among the most alternatively spliced transcripts in ELAVL1-depleted LSCs were regulators of mitochondrial function, providing the first link between not only RNA splicing, but a specific ELAVL1-mediated AS program and mitochondrial metabolism in LSCs. Furthermore, we identify for the first time TOMM34 as not only a key effector of ELAVL1 in human primary AML, but offering the possibility of mitochondrial import as a novel avenue through which LSC metabolism may be therapeutically targeted. TOMM34 is a co-chaperone that functions as the gateway mediating mitochondrial protein import through the translocase of outer mitochondrial (TOM) complex by stabilizing nuclear-encoded, mitochondrially destined pre-proteins in an unfolded state. Once shuttled across the TOM complex these highly diverse pre-proteins, which encode the majority of proteins necessary for mitochondrial function, then undergo more selective sorting and localization to specific membranes and regions within the mitochondria^107^. As such, targeting of mitochondrial import at the level of the ELAVL1-TOMM34 axis may serve as a more pervasive approach to disrupting localization and ultimately function, of a greater variety of mitochondrial proteins and thus offer the potential to solidify a robust inhibition of mitochondrial processes necessary for LSC maintenance and AML survival. Considered together, our findings here highlight a unique dependence of genotypically distinct primitive leukemic cells on oxidative metabolism that our data indicate can be counteracted by interfering with its post-transcriptional control via ELAVL1. Our work further provides insight into a clinically relevant connection between RBP-mediated post-transcriptional regulation and mitochondrial metabolism in AML LSCs that to our knowledge has not been elucidated to date.

Altogether our combined functional and molecular analyses showcase ELAVL1 as a critical novel regulator of LSCs that utilizes a combination of changes to the RNA landscape to coordinately enforce a state that supports LSC-optimal mitochondrial metabolism. Together with other diverse RBP regulators identified in our LSC-directed *in vivo* CRISPR screen, these findings highlight stem cell-adapted *in vivo* screening as a tractable tool to identify high value therapeutic targets, establish RBPs as essential players in LSC biology and open the door to elucidating and therapeutically exploiting their mechanisms of action.

## Acknowledgements

The authors thank Brad Doble, Bernardino Trigatti, Jon Draper and all members of the Hope lab for important feedback on this work; Minomi Subapanditha and Zoya Shapovalova for flow cytometry sorting; and Lillian Robson, Wendy Whittaker and Norma-Ann Kearns for animal caretaking and maintenance.

This work was supported by a Prins Bernhard Cultuurfonds Fellowship (LdR), a Canadian Cancer Society Post-doctoral Fellowship (SKL), a Canadian Institutes of Health Research (CIHR) MD/PhD Studentship (DC, TC), a Ontario Graduate Scholarship (DC, AV), a CIHR Doctoral Fellowship (AKC), a CIHR Canada Graduate Scholarship-Master’s Award (AT), a NIH R01 Grant (HL137223) (GWY, KH), an Ontario Institute for Cancer Research (OICR) Investigator Award (KH) and the OICR Acute Leukemia Translational Research Initiative (KH).

## Methods

### Mouse maintenance and transplants

B6.SJL (Ly5.1), C57Blk/6 (Ly5.2) and NSG mice (Jackson Laboratories) were bred and maintained in the Stem Cell Unit animal barrier facility at McMaster University. All procedures received the approval of the Animal Research Ethics Board at McMaster University. 24 h prior to transplantation by tail vein or intra-femoral injection, mice were sublethally irradiated (1 × 580 Rad). Bone marrow and spleen were harvested from moribund mice, crushed in RPMI + 10% FBS and passed through 40 uM cell strainers. Ammonium chloride was used for lysis of red blood cells.

### Leukemia and immortalized cell lines

RN2c cells (MLL-AF9/Nras^G12D^/hCas9; from Dr. Vakoc, Cold Spring Harbor Laboratory) were cultured in RPMI supplemented with 10% FBS, at a maximum density of 1 million cells per mL. MLL-AF9 and BC CML cell lines were generated as described in^29,57^ and cultured in SFEM (Stemcell Technologies) supplemented with 20 ng/μL SCF (Peprotech) and 10 ng/μL each of IL3 and IL6 (Peprotech). 293FT and HeLa (Invitrogen) cells were cultured in DMEM supplemented with 10% FBS. THP-1 (ATCC) cells were cultured in RPMI supplemented with 10% FBS. All cell lines tested negative for mycoplasma.

### Culture of primary AML patient samples

All AML patient samples were obtained with informed consent and with the approval of the local human subject research ethics board at University Health Network. Following ficol hypaque separation, mononuclear cells were stored in the vapor phase of liquid nitrogen in 10% DMSO, 40% FCS and alpha MEM. Primary samples were thawed in X-VIVO (Lonza) 50% FBS with 100 μg/ml DNAse prior to using in *in vitro* and *in vivo* assays. Primary AML samples were grown in AML growth media consisting of X-VIVO with 20% BIT Serum Substitute (Stem Cell Technologies) or StemSpan Serum Free Expansion Media II (SFEM II) (Stem Cell Technologies), supplemented with human stem cell factor (SCF, 100 ng/mL, R&D Systems), human interleukin-3 (IL-3, 10 ng/mL, R&D Systems), human interleukin-6 (IL-6, 20 ng/mL, Peprotech), human thrombopoietin (TPO, 20 ng/mL, Peprotech) and human FMS like tyrosine kinase 3 ligand (FLT3L, 100 ng/mL, R&D Systems).

### Lentiviral-infected and DHTS-treated primary AML transplantation assays

Production of shELAVL1 and shLuciferase/shScramble expressing lentiviral particles were performed as previously described^108^ and validated by RT-qpCR and/or western blot. For knockdown experiments, AML cells were thawed and transduced at an MOI of 50 for 24 or 48 hours, depending on the sample. For drug treatment experiments, AML cells were thawed and cultured with 1.1uM DHTS or equivalent volume of DMSO (vehicle). All cells were transplanted intrafemorally into sub-lethally irradiated (315 cGy) NSG mice at their corresponding timepoints. Mice were sacrificed 9 to 14 weeks post-transplant and bone marrow from the right femur (site of injection) and remaining tibias, pelvis and left femur were harvested along with spleens, filtered and red blood cell lysed using ammonium chloride (Stem Cell Technologies). Reconstituted mouse bone marrow and human AML was blocked with mouse Fc block (BD Biosciences) and human IgG (Sigma), respectively. Cells were subsequently stained with fluorochrome-conjugated antibodies: CD45 Pacific Blue, BV421; CD33 PE; CD14 PE-Cy7, APC-H7, CD11b BV605 and 7AAD PerCP-Cy5.5 for quantitative analysis by flow cytometry.

### Primary AML immunophenotyping and apoptosis

For knockdown experiments, AML cells were thawed, infected with lentivirus expressing pLKO.1-BFP-shScramble, -shELAVL1.1 or -shEALVL1.2 and cultured for 10 days. For DHTS experiments, AML cells were thawed and cultured with 5.4uM DHTS or equivalent DMSO volume for 48 hours. At their corresponding time points cells were blocked with human IgG (Sigma) and subsequently stained for quantitative flow cytometric analysis. For evaluation of knockdown experiments: CD14 PE; CD11b APC, 7AAD – PerCP-Cy5.5. For DHTS experiments: CD33 PE, BV605, CD14 FITC, APC-H7; CD11b BV605; Annexin V AlexaFluor 647.

### Isolation of human cord blood hematopoietic HSPCs

All patient samples were obtained with informed consent and with the approval of local human subject research ethics boards at McMaster University and the University Health Network. Human umbilical cord blood mononuclear cells were collected by centrifugation with Ficoll-Paque Plus (GE), followed red blood cell lysis with ammonium chloride (Stemcell Technologies). Cells were then incubated with a cocktail of lineage specific antibodies (CD2, CD3, CD11b, CD11c, CD14, CD16, CD19, CD24, CD56, CD61, CD66b, and GlyA; Stemcell Technologies) for negative selection of Lin^−^ cells using an EasySep immunomagnetic column (Stemcell Technologies). Live cells were discriminated on the basis of cell size, granularity and, as needed, absence of viability dye 7-AAD (BD Biosciences) uptake.

### AML and CB clonogenic progenitor assays

Thawed primary AML samples were counted and plated in a methylcellulose-based hematopoietic colony formation medium (Colony Gel, ReachBio), supplemented with DMSO or 1.1uM DHTS. Colonies were scored on days 10-14. Human CB samples were plated as described with a density of 1×10^3^ cells per 35mm plate. Cell suspensions were plated in duplicate and loose colonies consisting of 10 or more cells were scored and counted. THP-1 cells were plated at a density of 200.000 and 1000 cells per mL, respectively. Putative LSC-enriched populations were isolated from freshly expanded MLL-AF9 mouse leukemia cells by sorting c-Kit^high^ (top 10%) cells on a MoFlo XDP cell sorter (Beckman Coulter). Sorted cells were plated in triplicate in semi-solid methylcellulose medium (Methocult M3434; Stem Cell Technologies) at 1000 cells/mL. Colony counts were carried out after 10 days of incubation.

### sgRNA design and lentivirus production

sgNAs were designed using http://cripsr.mit.edu (quality score >70). For every gene, sgRNAs were targeted against RNA-binding domains (when annotated) or other Protein family (Pfam) domains to maximize negative selection phenotypes^30^ (see Supplementary Table 2 for an overview of all sgRNA sequences included in the libraries). sgRNAs were amplified as a pool and cloned into BsmBI digested pLKO1-CRISPR-H2B-GFP. Stbl4 electrocompetent cells (Invitrogen) were transformed, followed by DNA purification from 12 dishes of transformants (PureLink HIPure Plasmid Filter Maxiprep Kit, Invitrogen). Lentivirus was prepared by transient transfection of 293FT (Invitrogen) cells with pMD2.G and psPAX2 packaging plasmids (Addgene) to create VSV-G pseudotyped lentiviral particles. All viral preparations were titrated on HeLa cells before use on mouse bone marrow cells.

### *In vivo* pooled dropout screen transduction and transplantation

One million tertiary transplant RN2c cells per mouse were expanded *in vivo* (n= 4), leukemic bone marrow was harvested from moribund mice and cultured in fresh RPMI supplemented with 10% FBS and 5 μg/mL Polybrene (Sigma Aldrich). Pre-titrated lentivirus (RBP and NTC pools) was added at a clonal multiplicity of infection (MOI) of 0.2 with ∼300X coverage and cultures were incubated overnight after which the cultures were spun for 5 min at 1200 rpm, resuspended in fresh RPMI supplemented with 10% FBS and incubated for an additional 24 h. The H2B-GFP+ fraction was determined at 48 h post-transduction (18-22%), one-third of the cultures were frozen down for sequencing and remaining cells were transplanted into sublethally irradiated B6.SJL recipient mice (2 million cells per mouse, n= 7-9). After 10 days (T10 Primary), leukemic bone marrow and splenocytes were isolated and frozen down for sequencing and subsequent analysis (cells from every 2-3 mice were pooled to serve as biological replicates). Secondary mice were transplanted with a portion of the primary mouse bone marrow taken after thawing (T0 Primary) and at the 10-day endpoint (T10 Secondary) bone marrow and splenocytes again harvested (samples from every 2-3 mice were pooled). Total DNA from T0, T10 Primary, T0 Secondary and T10 Secondary cells was isolated using the DNeasy Blood & Tissue kit (Qiagen) according to the manufacturer’s instructions then further purified using RNeasy columns (69504, Qiagen). sgRNA sequences were then PCR-amplified using Q5 Hot Start High-Fidelity 2X Master Mix (M0494, NEB), barcoded primer pairs (see Supplemental Table 2) and 1 μg of DNA per PCR reaction. Individual 50 μL reactions were run on a 3% agarose gel and libraries were purified using the Zymoclean Gel DNA Recovery kit (D4007, Zymo Research). Sequencing was performed using standard Illumina instructions. MAGeCK^36^ (https://sourceforge.net/projects/mageck/) was used for analysis of sequencing reads and calculation of enrichment/depletion of individual sgRNAs. Sufficiency of sgRNA representation of greater than 500 was verified through all arms of the two-step screen.

### shRNA design and qRT-PCR

An Ametrine fluorescent protein was cloned into the pZIP-mCMV-ZsGreen-Puro vector (Transomic Technologies) by amplification of Ametrine from the MNDU3-MLL-AF9-PGK-Ametrine vector (kind gift from Dr.Guy Sauvageau) with addition of AclI/AgeI restriction sites and subsequent subcloning into AclI/AgeI digested pZIP. *Elavl1* shRNAs were adapted from the MISSION shRNA library (Sigma). Annealed oligos were digested with FastDigest XhoI/EcoRI (ThermoFisher) and cloned into the pZIP-mCMV vector. Confirmatory sequencing was carried out for all shRNAs before generation of lentivirus. Knockdown efficiency of shRNAs was determined by qRT-PCR. For all qRT-PCR determinations total cellular RNA was isolated with TRIzol LS reagent according to the manufacturer’s instructions (Invitrogen) and cDNA was synthesized using the qScript cDNA Synthesis Kit (Quanta Biosciences). The mRNA content of samples compared by qRT-PCR was normalized based on the amplification of GAPDH. qRT-PCR was done in triplicate with PerfeCTa qPCR SuperMix Low ROX (Quanta Biosciences) with gene specific probes (Universal Probe Library, UPL, Roche) and primers. See Supplemental Table 2 for all shRNA sequences.

### MLL-AF9 and bc-CML infections and transplants

MLL-AF9 or bc-CML cells were thawed and seeded in ultra-low attachment plates at a density of 0.5×10^6^ cells per mL of growth media. After 2-3 hours of recovery, cells were infected with pZIP-mCMV-Ametrine-shLuciferase, -shElavl1.6 or -shElavl1.7 at an MOI of 0.5 supplemented with 5 ug/mL polybrene at dose equivalents of ∼63, 000 and ∼31,000 cells per mouse. Cells were transplanted intravenously 24hr post-infection into sub-lethally irradiated (580 CyG) C57Blk/6 (Ly5.2) recipient mice. Bone marrow and spleens were harvested 14 and 9 days post-transplant for the MLL-AF9 and bc-CML recipients, respectively, for flow cytometric analysis.

### Isolation of mouse stem and progenitor cell populations

Bone marrow and spleen were harvested from 6-12 wk old mice, crushed in IMDM + 3% FBS and passed through 40 uM cell strainers. Ammonium chloride was used for lysis of red blood cells, followed by incubation of the cells with a cocktail of lineage specific antibodies (CD5, CD11b, CD19, B220, Gr-1 and TER119, Stemcell Technologies) for negative selection of Lin-cells using an EasyStep immunomagnetic column (Stemcell Technologies). Live cells were discriminated on the basis of cell size, granularity and, as needed, absence of viability dye 7-AAD (BD Biosciences) uptake.

### Competitive mouse HSPC transplants

Competitive transplants were performed according to^9^. Briefly, freshly sorted Lin-CD150+CD48-bone marrow cells were seeded at 3,000 cells per well in 96 well plates and cultured in SFEM supplemented with 10 ng/mL IL3 (Peprotech), 10 ng/mL IL6 (Peprotech), 100 ng/mL SCF (R&D) and 100 ng/mL TPO (Peprotech). Cultures were pre-stimulated for 24 h, followed by lentiviral infection with shRNAs targeting *Elavl1* and Luciferase (pZIP-mEF1a-ZsGreen-miR-E) in 8 μg/mL polybrene. Gene transfer (%) was determined 72 h post-transduction, followed by transplantation of the cultures (1/4 of each well per mouse, n = 4) along with 1×10^5^ whole bone marrow competitor cells (Ly5.2+). Donor-derived reconstitution was determined at 4-week intervals by examining peripheral blood samples from each mouse for Ly5.1+zsGreen+ fractions; BM and spleens were harvested from all recipients at 18 weeks post-transplant for immunophenotyping. For MLL-AF9 and BC CML cell lines, leukemic BM and splenocytes were freshly harvested from primary transplanted, moribund mice, followed by 24-48 h lentiviral infections with pre-titrated virus (pZIP-mCMV-Ametrine-shLuciferase/7sh*Elavl1*) in the presence of 5 μg/mL Polybrene. On the day of transplant, gene transfer % was determined by flow cytometry and 100.000 cells were intravenously transplanted into sublethally irradiated (580 cGy) C57Blk/6 recipient mice. The Ametrine+ fraction of the leukemic grafts was flow cytometrically analyzed from BM and spleens of moribund mice and compared to the Ametrine+ fraction at the day of transplant to determine the effect of KD on *in vivo* leukemic propagation.

### Human bone marrow cell subset RNA sequencing and analysis

Human adult bone marrow samples were collected from after obtaining informed consent according to procedures approved by the Research Ethics Board of the University Health Network. Hematopoietic stem and progenitors from bone marrow samples were sorted using a FACS Aria. 4-5 thousand HSC and progenitors were sorted from at least 3 bone marrow samples with the following phenotypes: LT-HSC, CD34+CD38-CD90+CD49f+; ST-HSC, CD34+CD38-CD90+CD49f-; MPP, multipotent progenitor (CD34+CD38-CD45RA-CD90-CD49f-). RNA from 2000 to 4000 cells was isolated using the PicoPure RNA isolation kit (Thermo Fisher). cDNA was generated and amplified using the SMART-SEQ2 method^109^. Sequencing libraries were generated with low with input RNA Nextera protocol (Illumina, Nextera DNA Sample Preparation Kit, Cat No. FC-121-1031). The samples were sequenced on Illumina HiSeq2500 with V4 flowcells. RNA-seq data was mapped using STAR aligner^110^. The FPKM and expression values were calculated using Tuxedo suits tool kit^111^ and R was used for downstream statistical analysis.

### Enhanced Cross-linking and Immunoprecipitation (eCLIP)

BC CML cells were freshly harvested, crushed in IMDM + 3% FBS, passed through 40 uM cell strainers followed red blood cell lysis. Cells were incubated with a cocktail of lineage specific antibodies (CD5, CD11b, CD19, B220, Gr-1 and TER119, Stemcell Technologies) for negative selection of Lin-cells using an EasyStep immunomagnetic column (Stemcell Technologies). 20 million cells were per sample were subsequently washed in PBS and UV-crosslinked at 400mJ/cm^2^ on ice. Samples were then pelleted, snap-frozen and stored at -80°C. eCLIP-seq was performed as previously described^63^. Pellets were lysed in iCLIP lysis buffer, treated with RNAse I for 5 min at 37°C, followed by immunoprecipitation using 10 μg anti-ELAVL1 mouse monoclonal IgG (ab170193, Abcam) and 125 μL M-280 Sheep-α-Mouse IgG Dynabeads (ThermoFisher #11201D) per sample. After stringent rounds of washing, samples were dephosphorylated (FastAP; ThermoFisher, T4 PNK; NEB), 3’ ligation (on-bead) with a barcoded RNA adapter followed using T4 RNA Ligase (NEB). Samples were again stringently washed, run on standard protein gels and transferred to nitrocellulose membranes. The region spanning 36-115 kDa was then isolated, followed by extraction of RNA from the membranes and reverse transcription (AffinityScript; Agilent). A DNA adapter containing a 5’ random-mer was then ligated (3’), followed by cleanup of the samples and PCR amplification. Libraries were size-selected (175-350 bp) and purified from 3% low melting temperature agarose gels, followed by sequencing on the Illumina HiSeq 2500 platform (paired-end). A novel eCLIP processing pipeline (Yeo lab in-house generated) was used for analysis of the sequencing reads, followed by downstream analysis using BEDTools, HOMER and Pathway Analysis. See Supplementary Table 2 for RNA and DNA adapters used. Removing potential PCR duplicates revealed 3,532,702 and 4,027,997 usable reads in the two ELAVL1 immunoprecipitation (IP) replicates. Corresponding size-matched input controls found 1,525,778 and 1,394,954 usable reads. After identifying peaks enriched above input control (-log10(p-value) ≥ 3 and log2(fold) ≥ 3) and irreproducibility discovery rate analysis^63^.

### RNA-sequencing of RN2c cells

RN2c cells were transduced with lentiviral-packaged sgRNA targeting ELAVL1 and, as control, Ano9. 48 hours following transduction live 7AAD-GFP+ cells were isolated by FACS and RNA was extracted by TRIzolLS (ThermoFisher # 10296028) following manufacturers protocol. RNA was coprecipitated with GlycoBlue (ThermoFisher #AM9516) following manufacturers protocol; in brief, 3uL (final concentration 45ug/mL) of GlycoBlue was added to TrizolLS-separated aqueous phase followed by one volume isopropanol and incubated at -20°C for 30 min before pelleting RNA by centrifugation. Isolated RNA was treated with DNase I (ThermoFisher #EN0521) which was inactivated by addition of EDTA and incubation at 65°C. PolyA+ RNA libraries were generated using NEBNext PolyA mRNA Magnetic Isolation Module (E7490) and NEBNext Ultra II Directional RNA Library Prep Kit for Illumina (E7760L) and indexed using NEBNext Multiplex Oligos for Illumina (E6440L). 100bp paired-end sequencing was performed on the Illumina HiSeq 1500 at a depth of 50 million reads per sample. Sequencing reads were processed to identify differential gene expression via Yeo lab in-house pipeline that makes use of the fastQC (v0.10.1), cutadapt (v1.14.0), STAR (v2.4.0), featureCounts (v1.5.0-p1) and DESeq2 (v1.14.1) packages. Alternative splicing was determined using rMATS (v3.2.5) and significant non-overlapping alternative splice events was assigned where FDR <0.05 and lncLevelDifference was magnitude 0.1 or higher.

### RNA-sequencing of primary AML cells

Human primary AML cells (AML #4) were infected in triplicate with pLKO.1-EGFP-shScramble or - shELAVL1.7 at an MOI of 50 for 48hr after which 7AAD^-^EGFP^+^ cells were sorted and 100, 000 cells per condition were resuspended in TRIzolLS (ThermoFisher Scientific; catalog #10296028) following manufacturer’s protocol. Aqueous phase was separated using Phasemaker Tubes (ThermoFisher Scientific; catalog #A33248) and RNA was isolated using TRIzolLS manual both following manufacturer’s protocol. Isolated RNA was treated with DNase from TURBO DNA-free kit (ThermoFisher Scientific; catalog #AM1907) following manufacturer’s protocol. 100bp paired-end sequencing was performed on the Illumina HiSeq 1500 at a depth of 50 million reads per sample. Sequencing reads were aligned to the hg38 reference genome using STAR (v2.7.2c). Quantification and differential expression was performed using RSEM (v1.3.1) and DESeq2 (v1.26.0). Alternative splicing was determined using rMATS (v4.0.2) and significant non-overlapping alternative splice events were assigned where FDR <0.05 and lncLevelDifference was magnitude 0.1 or higher.

### Expression profiling analysis of AML patient diagnosis-relapse paired samples

ELAVL1 expression values were normalized in logTPM from three publicly available datasets^3,54,55^ and a Wilcoxon signed-rank test was used to compare the groups. We included only pairs in which the relapse followed chemotherapy treatment.

### Gene Set and Pathway Analysis

GO annotations were assigned to screening candidates and hits using the open-source web-application GOnet (Pomaznoy M, Ha B, Peters B. GOnet: a tool for interactive Gene Ontology analysis. BMC Bioinformatics. 2018;19:470. Above and below-median cohorts for ELAVL1 expression were identified in the BeatAML RNA-seq dataset^74^ using DESeq-normalized counts, and altered genesets were identified using DESeq2 followed by fGSEA. The fgsea R package (v1.12.0) was used to perform preranked GSEA against a gene set repository maintained by the Gary Bader Lab, which encompasses gene sets from GO Biological Processes, Reactome, and MSigDB. Rank scores were calculated as -log10(p)*sign(log2FC). Enrichments were visualized using the Cytoscape Enrichment Map plugin. Pathway analyses for differentially spliced genes and eCLIP targets were performed using g:Profiler.

### Flow cytometry

All flow cytometry analysis was performed using a MACSQuant Analyzer 10 (Miltenyi Biotec), BD LSRII Analyzer or BD LSR Fortessa X-20, analysis was performed using FlowJo software (Tree Star).

### Intracellular flow cytometry

Primary AML cells infected with lentivirus expressing pLKO.1-BFP-shScramble or -shELAVL1 were fixed and permeabilized for 20min at 4C using BD Cytofix/Cytoperm Fixation/Permeabilization Solution Kit 48hr post-infection. Cells were subsequently incubated individually with mouse anti-HuR (Abcam; 1:500) or mouse anti-c-myc (Abcam; 1:500) overnight at 4C. Cells were washed and stained with AlexFluor 647 donkey anti-mouse (1:2500) for 1hr at 4C and subsequently analyzed by flow cytometry.

### MitoTracker experiments

Primary AML cells were incubated with 2uM FCCP (Cedarlane) for 30min at 37C followed by 2uM FCCP with 50nM MitoTracker Orange CMTMRos (MTO) (ThermoFisher) for 30min at 37°C 72hr post-infection. RN2c cells were incubated with 100nM MTO for 30min at 37C 48hr post-infection. Primary AML cells were stained with 7AAD and RN2c cells were stained with SytoxBlue at their corresponding timepoints. The median florescence intensity of MTO in live cells was quantified by flow cytometry.

### Western blot

Immunoblotting was performed with anti-ELAVL1 mouse monoclonal IgG (ab170193, Abcam), anti-RBM14 rabbit polyclonal (ab70636, Abcam) and α-Tubulin rabbit monoclonal IgG (2125S, Cell Signaling Technologies) antibodies. Secondary antibodies used were IRDye 680 goat anti-rabbit IgG and IRDye 800 goat anti-mouse IgG (LI-COR).

### Immunofluorescence

Approximately 180.000 primary AML or RN2c cells were prepared for staining by cytocentrifugation (600 rpm, 5 min). Cells were fixed in 4% PFA for 20 minutes, followed by permeabilization in 0.1X Triton X-100 in PBS for 10 min. Samples were incubated in blocking buffer (PBST + 10% Goat serum + 1% W/V BSA) for 1 hr at room temperature, followed by incubation with anti-ELAVL1 (ab170193, Abcam) and anti-G3BP1 (13057-2-AP, Proteintech), overnight at 4°C. Secondary antibody incubation was performed with AF647 donkey-anti-mouse IgG or AF488 donkey-anti-rabbit IgG for 1 hr at room temperature, DAPI staining was performed simultaneously. Slides were mounted with Fluoromount mounting medium, images were captured using an Olympus IX81 microscope (40X objective lens).

### Statistical analysis

Unless stated otherwise (i.e., analysis of RNA-seq data sets), all statistical analysis was performed using GraphPad Prism (GraphPad Software version 5.0). Unpaired student t-tests were performed with p < 0.05 as the cut-off for statistical significance. CRISPR scores from ^40^ were analyzed using MATLAB (R2014B, The MathWorks, Inc software).

