## Supplemental Figures for "A two-step *in vivo* CRISPR screen unveils pervasive RNA binding protein dependencies for leukemic stem cells and identifies ELAVL1 as a therapeutic target"

### Supplemental Figure 1

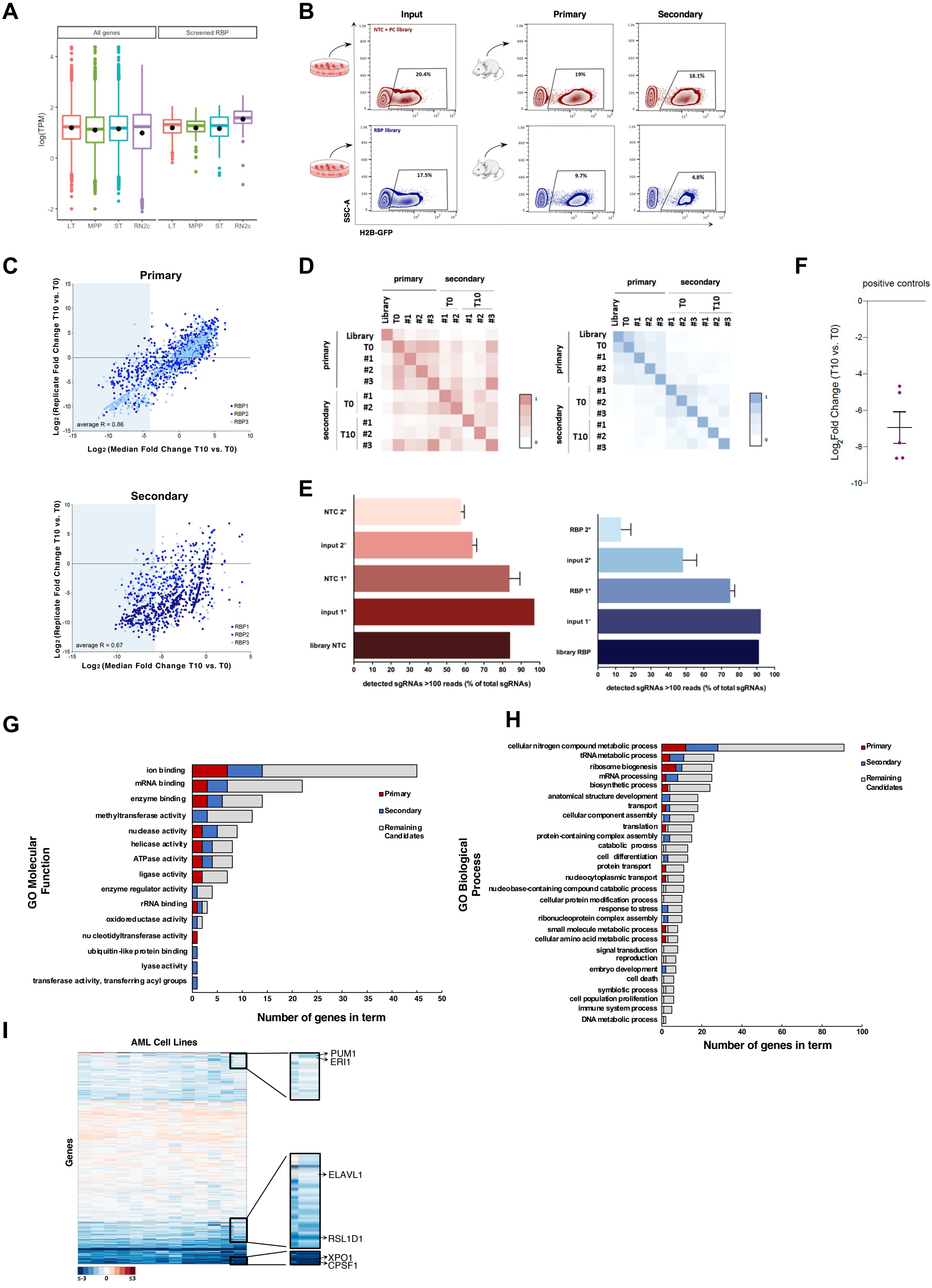

**Figure S1. Two-step *in vivo* CRISPR-Cas9 screening of LSC-associated RBPs identifies drivers of AML LSC function.** (A) Gene expression for all genes in comparison to the screened RBPs in control RN2c cells vs purified mouse bone marrow stem and progenitor cell fractions from[112]. (B) H2B-GFP$^{+}$ fractions of input RN2c cultures 48hr post-transduction (T0) and representative Ly5.2$^{+}$ endpoint grafts (T10) from the primary and secondary transplants. (C) sgRNA dropout reproducibility across duplicate RBP-targeting primary (top) and secondary (bottom) arms of the screen; scatter plot illustrates the correlation of normalized reads per sgRNA in three independent replicates. Blue shaded area represents sgRNAs depleting >20 fold (on average). R = pearson correlation coefficient. (D) Pearson correlation coefficients for the normalized sgRNA read counts of the plasmid library in cells 48hr after lentiviral infection (T0) and BM samples 10 days post-transplant (T10) for both primary and secondary rounds of the NTC (left) and RBP (right) arms of the screen. T0 in the secondary arm indicates the fraction of primary transplant endpoint samples that was retained and library representation at the endpoint of secondary grafts (secondary T10) was compared to this input. (E) Percentage of all sgRNAs at each sampling point in the NTC (left) and RBP (right) arms of the screen detected at a read count great than 100. (F) $Log_2$ fold change (LFC) of positive controls targeting sgRNAs in the primary arm of the dropout screen. (G) GO annotation plots illustrating associated Biological Processes (left) and Molecular Functions (right) for hits and RBP screen candidates. (H) CRISPR score (average $log_2$ fold-change of sgRNAs after 14 *in vitro* population doublings) of >18.000 genes (rows) in 13 human AML cell lines [40] (columns). A low CRISPR score corresponds to a high degree of essentiality. Arrows indicate CRISPR scores of select primary and secondary RN2c screen hits.

### Supplemental Figure 2

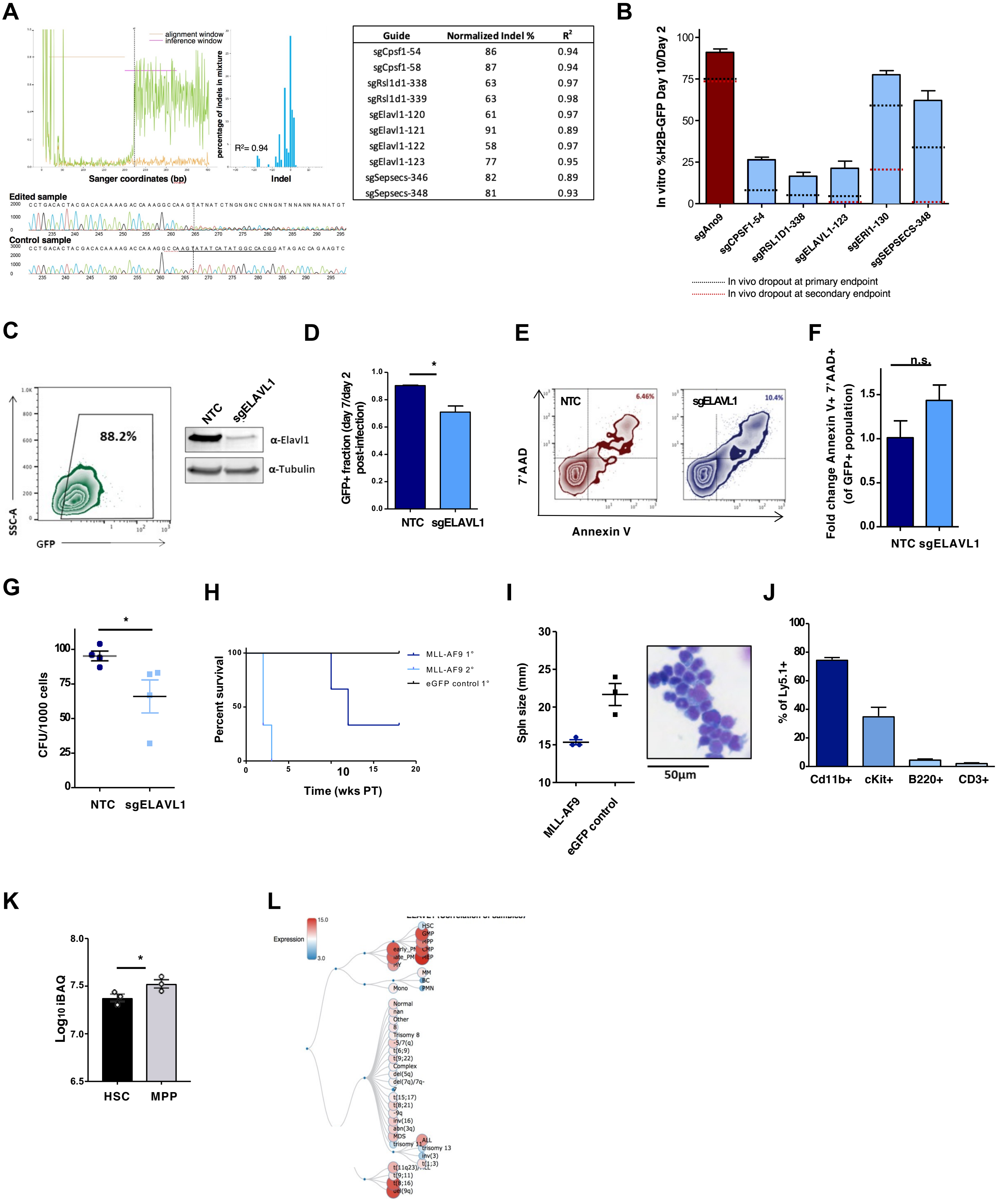


| Guide | Normalized Indel % | $R^2$ |
|---|---|---|
| sgCpsf1-54 | 86 | 0.94 |
| sgCpsf1-58 | 87 | 0.94 |
| sgRsl1d1-338 | 63 | 0.97 |
| sgRsl1d1-339 | 63 | 0.98 |
| sgElavl1-120 | 61 | 0.97 |
| sgElavl1-121 | 91 | 0.89 |
| sgElavl1-122 | 58 | 0.97 |
| sgElavl1-123 | 77 | 0.95 |
| sgSepsecs-346 | 82 | 0.89 |
| sgSepsecs-348 | 81 | 0.93 |

**Figure S2. Validation and characterization of primary and secondary screen hit-targeting sgRNAs.** (A) Representative indel plots and traces generated from ICE analysis[41] for individually validated sgRNAs in test RN2c cells. Level of discordance (top left) in the sample sequence (green) and control sequence (orange) before and after the expected cut-site is shown. Indel plot (top right) shows percentages of the various calculated indel sizes. Trace sequences (bottom) display the aligned sample and control sequences with the targeted cut-site. Indel percentages for individually validated sgRNAs normalized to the percent infection of each sample. $R^2$ values were generated from Indel plots. (B) *In vitro* assessment of RN2c growth upon knockout of selected screen hits. Levels of H2B-GFP reduction achieved in parallel *in vivo* experiments in primary and secondary mice are highlighted in black and red dotted lines respectively. (C) Western blot validation of ELAVL1 knockout by pL.CRISPR.EFS.GFP-sgELAVL1 in HEK293 cells one week post-infection. Prior to lysis, cultures were analyzed by flow cytometry for the fraction of $GFP^+$ cells; sgNTC was used as a negative control. (D) sgRNA-mediated KO of ELAVL1 in THP-1 cells; $GFP^+$ fraction was followed over time. $GFP^+$ fractions of sgNTC and sgELAVL1 infected THP-1 cells at 7 days post-transduction are normalized to the percent infected at 2 days PT. Representative flow plots (E) and quantitative analysis (F) of the apoptotic fraction of $GFP^+$ sgNTC and sgELAVL1 THP-1 cells 4 days post-plating. Data represents 3 replicate infections. (G) CFU assays performed with FACS-purified $GFP^+$ sgNTC and sgELAVL1 THP-1 cells. (H) B6.SJL mouse LSK cells were retrovirally infected with MSCV-MLLAF9-PGK-eGFP (or MSCV-PGK-eGFP, control) and GFP+ cells FACS-purified 48hr post-infection transplanted into C57Blk/6 recipients. Fully engrafted primary MLL-AF9-driven leukemic BM was challenged in secondary recipients. Latency of primary (1°) and secondary (2°) transplant MLL-AF9 leukemias, as well as primary eGFP control BM (n = 3 for each arm) is shown. (I) Spleen sizes of primary engrafted MLL-AF9-driven leukemias and primary eGFP control BM (left) and Wright-Giemsa staining of peripheral blood sampled from primary engrafted recipient mouse in MLL-AF9 arm (right). (J) Flow cytometric evaluation of the immunophenotype of MLL-AF9-driven BM grafts. (K) Expression levels of Elavl1 in mouse HSC (LSK CD34-CD150+CD48-FLK2-) and MPP (LSK CD34+CD15-+CD48-FLK2-) populations[42]. Intensity based absolute quantification (iBAQ) levels are shown. (L) Correlation tree of normal and malignant hematopoietic samples based on *ELAVL1* expression levels generated by the BloodPool tool, adapted from Bloodspot[43] showcasing low (blue) expression in HSCs relative to higher (pink to red) expression in bulk AML cells.

### Supplemental Figure 3

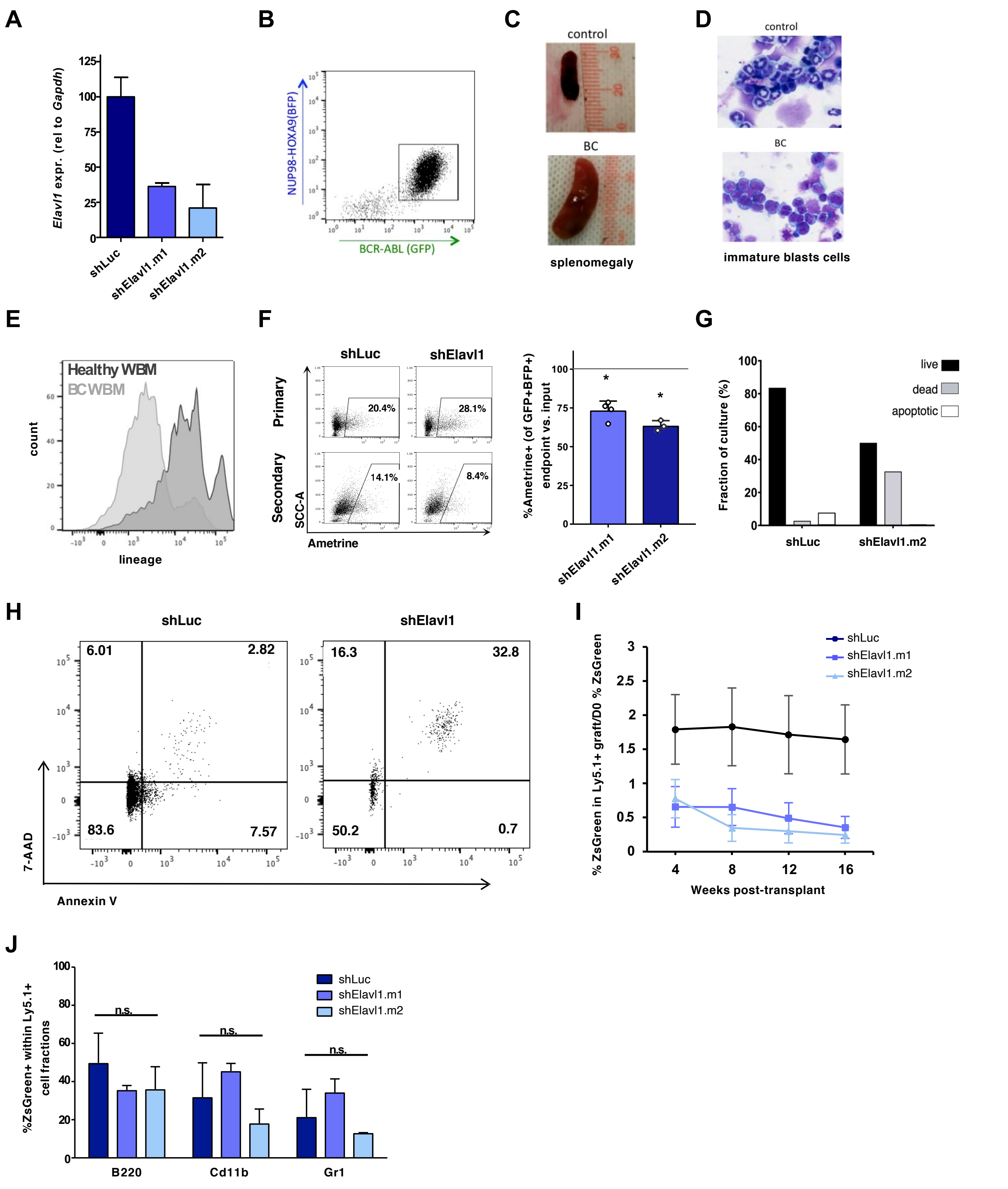

**Figure S3. Generation of mouse myeloid leukemia models and analysis of Elavl1 knockdown in murine healthy versus malignant hematopoietic cells.** (A) qPCR knockdown validation of shElavl1.6 and shElavl1.7 Ametrine$^+$ cells in primary mouse BM. Data is normalized against *Gapdh*. (B) B6.SJL LSK cells were retrovirally infected with NUP98-HOXA9-BFP and BCR-ABL-GFP. BFP$^+$GFP$^+$ cells isolated 48h post-infection were transplanted into C57Blk/6 recipients. Flow cytometric evaluation of endpoint blast-crisis BM grafts (10 days post-transplantation) is shown. (C, D) Splenomegaly (C) and infiltration of BM with immature blast cells (Wright-Giemsa staining of peripheral blood sampled from primary engrafted recipient mouse) (D) indicate advanced stages of disease (blast-crisis) 10 days PT. (E) Flow cytometric evaluation of recipient mice whole BM (WBM) at blast-crisis shows a drastic decrease in lineage positive cells as compared to WBM from healthy mice (control). *$p < 0.05$ as determined by a two-sided Student's t test. (F) Output vs. input analysis of the Ametrine$^+$ fraction of shElavl1 transduced BC CML BM. GFP$^+$BFP$^+$ BM was analyzed at the endpoint of secondary transplantation; data is normalized to shLuciferase (dotted line). Representative flow plots are shown at left. (G) Fraction of apoptotic cells was measured in shLuciferase- and shElavl1-transduced BC CML cultures, 3 days post-infection. Flow plots are shown in (H). (I) Percentage of ZsGreen$^+$ cells within Ly5.1$^+$ fractions in peripheral blood (PB) samples at 4-week intervals post-transplant relative to the percent of ZsGreen$^+$ cells at day 0 (D0). (J) Quantification of lineage marker expression in 18-week endpoint BM grafts initiated by shLuciferase, shElavl1.6- or shElavl1.7-transduced cells.

### Supplemental Figure 4

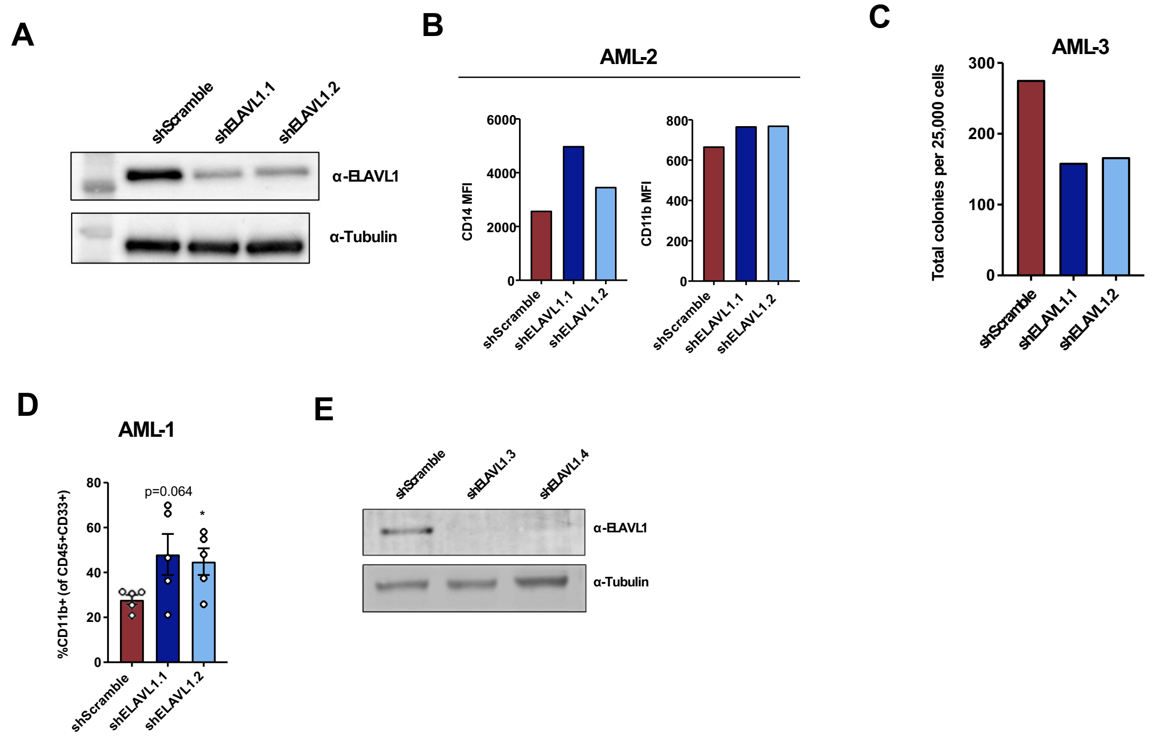

**Figure S4. ELAVL1 knockdown promotes AML cell differentiation.** (A) Western blot validation of ELAVL1 knockdown by shELAVL1.1 and shELAVL1.2 in HeLa cells, normalized to α-tubulin. (B) Median fluorescence intensities (MFI) of CD14 and CD11b in primary AML cultures 10 days post-infection. (C) CFU output from infected primary AML cells 16 days post-plating (n=1). (D) Flow cytometric analysis of $CD11b^+$ populations in bone marrow at endpoints from shScramble- and shELAVL1.1-infected grafts. (E) Western blot validation of ELAVL1 knockdown by shElavl1.3 and shElavl1.4 in HEK293 cells, normalized to α-tubulin.

### Supplemental Figure 5

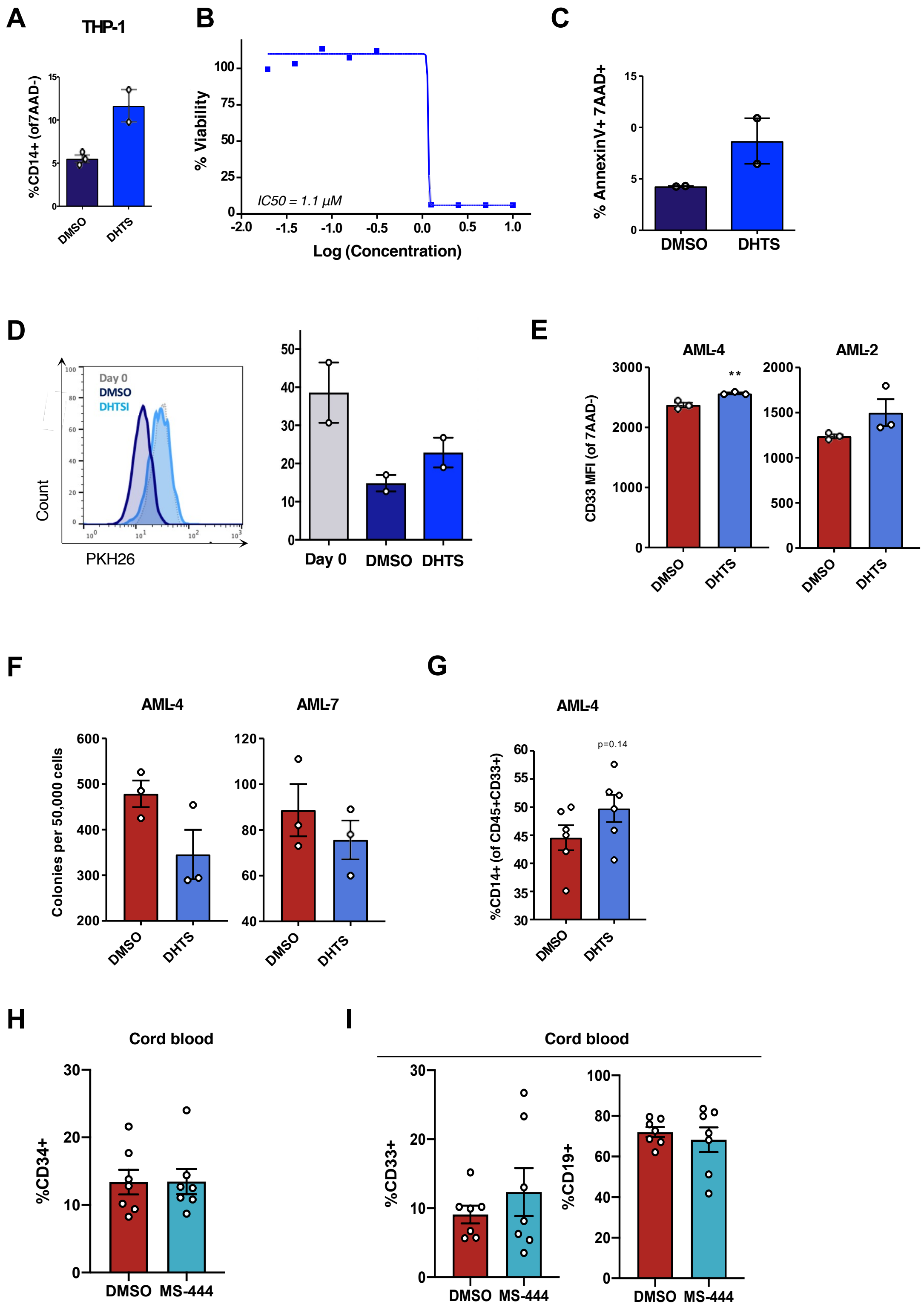

**Figure S5. ELAVL1 inhibitors restrict proliferation and promote differentiation of AML cells. (A)** Flow cytometric evaluation of the $CD14^+$ fraction of THP-1 cells 96 hours post DMSO or DHTS (1.1uM) treatment. (B) IC50 curve of DHTS-mediated inhibition of THP-1 cells 72hr post-treatment. (C) Flow cytometric analysis of the apoptotic fraction ($AnnexinV^+7AAD^+$) of DMSO- and DHTS-treated (1.1 μM) THP-1 cells 96 hours post-treatment. (D) PKH26 labeling of DMSO- and DHTS (1.1 μM) treated THP-1 cells before treatment (day 0) and 48 hours post-treatment. n = 2 replicate experiments. (E) Median fluorescence intensity of CD33 in human primary AML 48hr post-DMSO or -DHTS (5.4uM) treatment. n = 3 replicate experiments. (F) CFU output from DMSO- or DHTS-treated (1.1uM) human primary AML samples. (G) Flow cytometric analysis of injected femur from DMSO- and DHTS-treated human primary AML recipient mice showing the $\%CD14^+$ fraction in the leukemic graft. (H, I) Flow cytometric analysis of HSPC (CD34+) (I) and lineage (myeloid – CD33; lymphoid – CD19) markers (J) in CB grafts from DMSO- and MS-444-treated mouse bone marrow. n.s = not significant, *p < 0.05, **p < 0.01, determined by a two-sided Student's t test.

#### Supplemental Figure 6

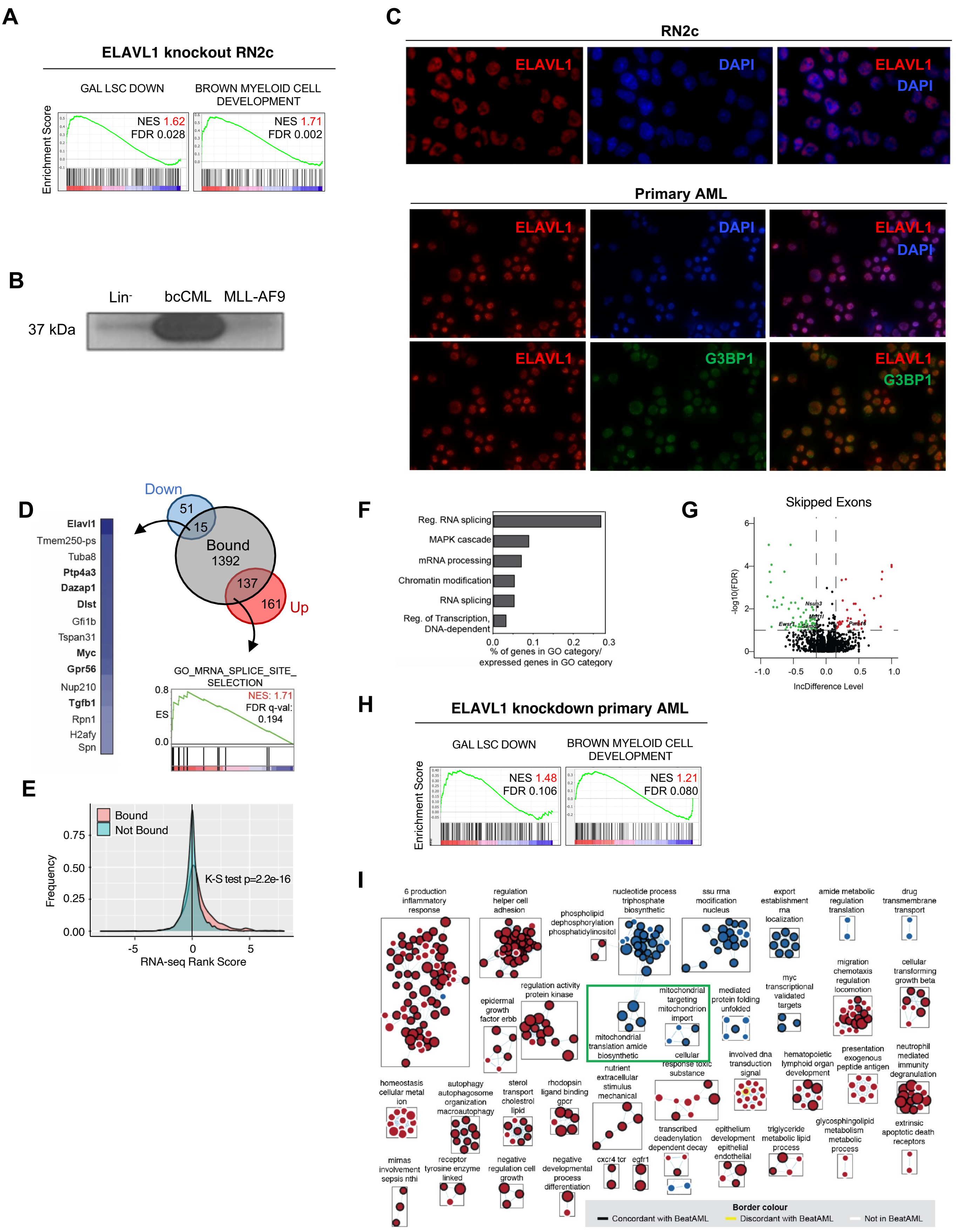

**Figure S6: Characterization of ELAVL1 targets and global influence on the transcriptional landscape.**

(A) GSEA plots showing LSC and myeloid signature enrichments in the Elavl1-knockout RN2c transcriptome. (B) Western blot of Elavl1 in murine Lin$^-$, bcCML and MLL-AF9 cells. (C) Immunofluorescent microscopy of ELAVL1 and nuclear marker (DAPI) and cytoplasmic marker (G3BP1). (D) Summary of intersection of transcripts bound (grey) and differentially expressed (up=red, down=blue, p. adj <0.05) upon Elavl1-deletion. (E) Expression distribution of bound and not bound transcripts in Elavl1-knockout RN2c cells. (F) GO term enrichment analysis for transcripts both bound and differentially expressed (p adj <0.05) upon Elavl1-deletion. (G) Volcano plot of differential exon skipping in ELAVL1 knockout RN2c cells, >15% inclusion and exclusion at FDR<0.1 indicated in red and green respectively. (H) GSEA plots of LSC and myeloid signature enrichments in AML. (I) Enrichment map of pathways significantly (FDR<0.1) changed in human AML following ELAVL1 knockdown with borders demarcating concordance/discordance with enrichments from below-median ELAVL1-expressing samples from the BeatAML data set**.**

#### Supplemental Figure 7

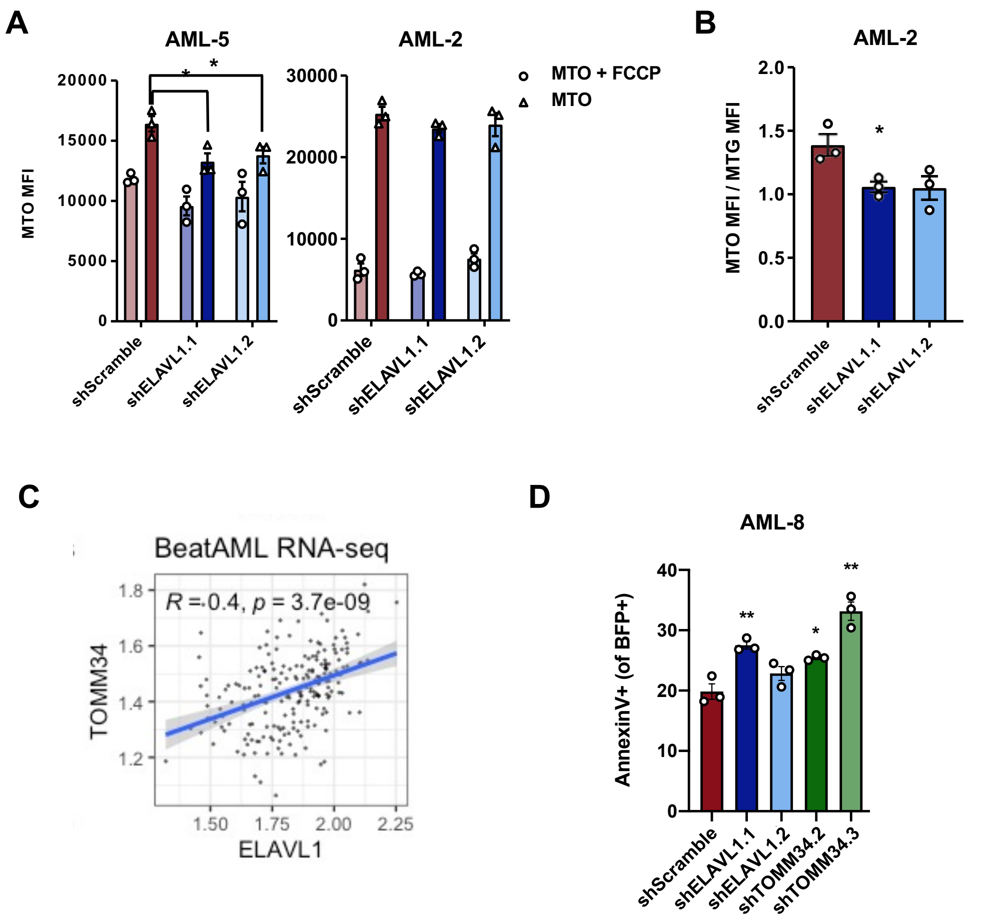

**Figure S7: TOMM34 depletion in human primary AML phenocopies ELAVL1 and impairs mitochondrial function.** (A) Quantification of MTO median fluorescence intensity in the presence and absence of FCCP in human AML. (B) Membrane potential as measured by the fraction of active mitochondria (MTO MFI) from total mitochondrial mass (MitoTracker Green (MTG)) in human AML upon ELAVL1 loss. (C) Correlation of TOMM34 and ELAVL1 expression in the BeatAML dataset. (D) Apoptotic analysis (AnnexinV+) of ELAVL1- and TOMM34-depleted human primary AML cultures.
